# Methylene blue modulates G3BP1-associated condensation and attenuates *Kras*-driven pancreatic preneoplastic lesions

**DOI:** 10.64898/2026.11.09.765332

**Authors:** Zhongjun Zhao, Jing Xue, Wenhui Zhang, Quanda Li, Lele Cheng, Xiaoya Wu, Yifu Zhu, Xue Luan, Yan Huang, Na Wang, Liye Ma, Jintao Xu, Ronghua Yu, Jingke Ding, Chong Hong, Huixia Zhang, Yuran Zhao, Chunli Yong, Mingyong Liao, Xuan Shi, Quanlin Zhao, Yanru Chen, Shuming Zhou, Xinghua Huang, Christian Wolfrum, Tongtong Wang, Wenjie Ge

## Abstract

Stress granules (SGs) are dynamic biomolecular condensates implicated in pancreatic ductal adenocarcinoma (PDAC), with G3BP1/2 serving as key regulators of their assembly. We combined structure-based screening of FDA-approved drugs with cellular and biochemical assays and identified methylene blue (MB) as a modulator of SG assembly. MB suppressed SG accumulation across multiple PDAC cell lines under arsenite (AS)-induced stress, nutrient deprivation, and heat shock without reducing the abundance of major SG-associated proteins. In reconstituted systems, MB inhibited G3BP1 condensation with or without added RNA and showed a nonsignificant trend toward lower G3BP1–Caprin1 co-immunoprecipitation in cells. Molecular docking and molecular dynamics simulations predicted a potential MB interaction site within the G3BP1 NTF2-like (NTF2L) domain. The G3BP1 V11A/F15A/F33A/F124A mutant retained basal condensate-forming capacity but showed reduced responsiveness to MB compared with wild-type G3BP1, implicating this region in MB-mediated regulation. In *Kras*-driven KC mice, chronic oral MB administration was associated with reduced pancreatic G3BP1-positive puncta, increased acinar preservation, and decreased preneoplastic lesion burden, epithelial proliferation, and stromal remodeling, without overt abnormalities in the evaluated endpoints. Together, these findings identify MB as a small-molecule probe that modulates G3BP1 condensation dynamics and demonstrate that chronic MB treatment is associated with the attenuation of early *Kras*-driven pancreatic preneoplastic lesion progression in vivo.

## Introduction

Stress granules (SGs) are dynamic, membrane-less ribonucleoprotein condensates that assemble in response to cellular stress through multivalent interactions and liquid– liquid phase separation (LLPS) (1–3). By organizing untranslated messenger ribonucleoproteins (mRNPs) and RNA-binding proteins (RBPs), SGs contribute to cellular stress adaptation, whereas their dysregulation has been implicated in cancer and neurodegenerative diseases (4, 5). In pancreatic ductal adenocarcinoma (PDAC), nutrient deprivation, hypoxia, and inflammatory stress may promote SG assembly and contribute to tumor cell survival and therapeutic resistance (6–10). Recent studies have further implicated SGs in PDAC cell-state heterogeneity, therapeutic responses, and early *Kras*-driven pancreatic lesion development (9, 11).

G3BP1 and G3BP2 are central regulators of SG assembly (12–14). Their N-terminal nuclear transport factor 2-like (NTF2L) domain plays a dual structural role: it forms a constitutive homodimer scaffold that provides basal valency while presenting a distinct, highly conserved hydrophobic pocket on each protomer that recognizes short linear motifs (such as FGDF and FxFG motifs) in regulatory partners, including Caprin1 and USP10 (12–18). While the dimer core remains structurally stable, this peripheral interaction groove dictates the functional balance between condensation-promoting and -inhibiting complexes, rendering this pocket an attractive site for small-molecule intervention without necessarily perturbing the underlying homodimeric framework.

Previous studies have explored peptides and small molecules targeting G3BP1-associated interactions (16, 19), but their development as selective pharmacological modulators remains at an early stage. Drug repurposing offers an alternative strategy for identifying SG modulators from compounds with established pharmacological and safety profiles (20).

Here, we combined structure-based screening of 2,783 FDA-approved compounds with cellular and biochemical assays and identified methylene blue (MB) as a modulator of G3BP1-associated condensation. We examined its effects on SG assembly under different stress conditions, G3BP1 condensation in reconstituted systems with and without added RNA, and G3BP1 – Caprin1 association. Computational and mutational analyses were used to investigate the potential involvement of the NTF2L interaction surface. Finally, we evaluated the effects of MB on SG-associated changes and pancreatic preneoplastic lesions in *Kras*-driven mice.

## Results

### Structure-based screening of an FDA-approved drug library identifies four compounds that suppress stress granule accumulation

We performed structure-based virtual screening of 2,783 compounds from the FDA-Approved Drug Library (MedChemExpress, MCE) against the G3BP1 NTF2L domain, followed by cell-based screening for compounds that reduce SG accumulation (Fig. 1A; Fig. S1A–B). Successive screening and validation in CFPAC-1 cells identified four candidates: homoharringtonine (HY-14944), fedratinib hydrochloride hydrate (HY-10409A), sanguinarine chloride (HY-N0052A), and methylene blue (MB; HY-B1359) (Fig. 1B–C). Their docking scores and predicted binding poses are presented in Fig. S1B–C. All four compounds reduced sodium arsenite (AS)-induced G3BP1-positive puncta, with MB showing a pronounced effect (Fig. 1D).

**Fig. 1.**
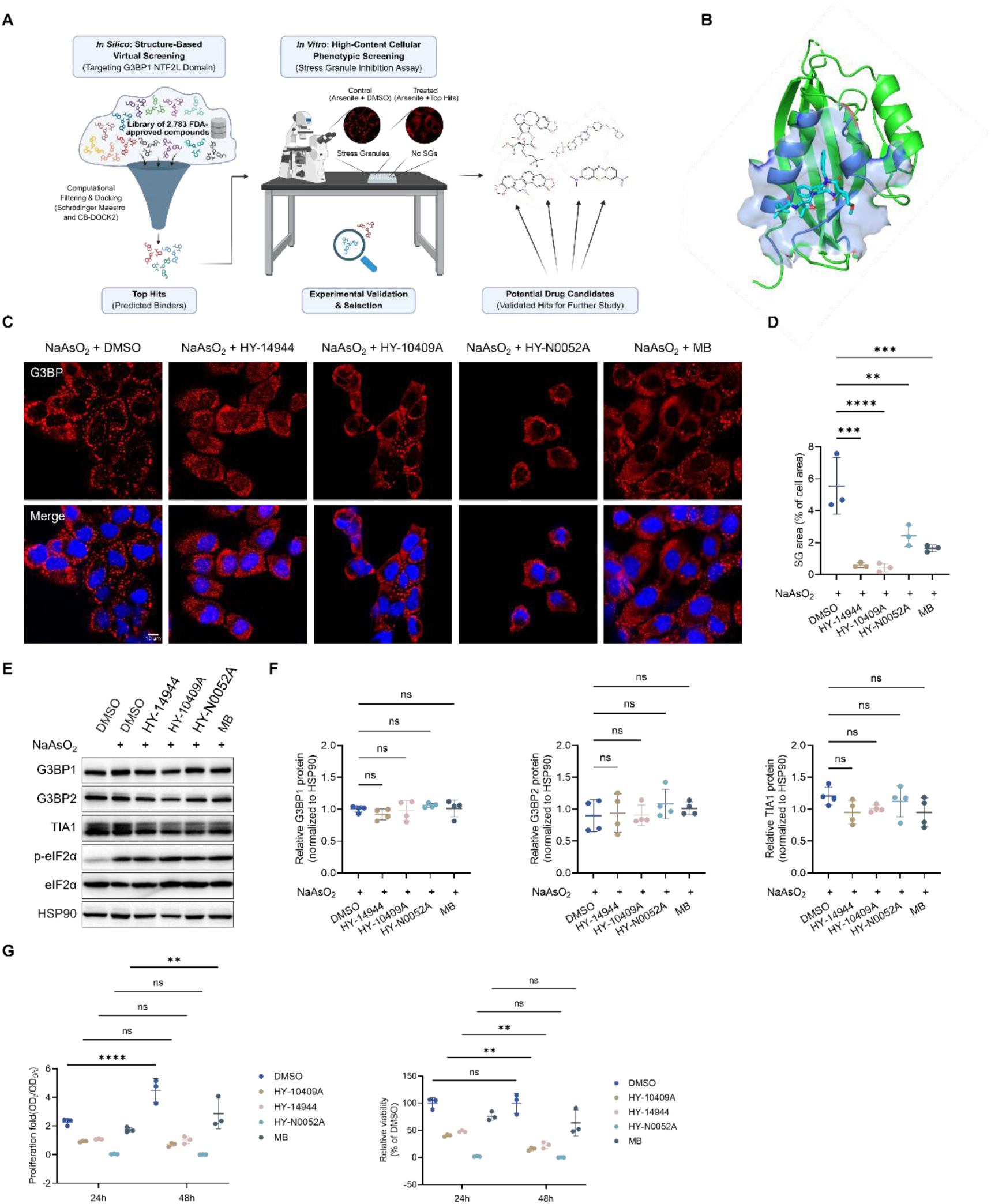
Structure-based screening of an FDA-approved drug library identifies four compounds that suppress stress granule accumulation. (A) Schematic overview of the docking-guided compound screening and cellular validation workflow. (B) Structural visualization of the predicted ligand-binding pocket within the NTF2-like domain of G3BP1, showing the docking poses of candidate compounds. G3BP1 is displayed in cartoon representation, the binding pocket as a semitransparent molecular surface, and the docked ligands as stick models. (C) Representative IF images of G3BP1-positive puncta in CFPAC-1 cells treated with AS (1 mM) and the indicated compounds (10 μM) for 1 h. Scale bar, 10 μm. (D) Quantification of SG accumulation, expressed as the percentage of cellular area occupied by G3BP1-positive puncta. Data are presented as mean ± SD from three independent biological experiments (n = 3). Statistical significance was determined using ordinary one-way ANOVA followed by Dunnett’s multiple-comparisons test against the AS-treated vehicle control. (E) Representative immunoblots showing G3BP1, G3BP2, and TIA1, phosphorylated eIF2α (p-eIF2α), and total eIF2α following treatment with the indicated compounds. HSP90 served as the loading control. (F) Densitometric quantification of G3BP1, G3BP2, and TIA1 protein abundance in (E), normalized to HSP90. Data are presented as mean ± SD (n = 4). Statistical significance was determined using ordinary one-way ANOVA followed by Dunnett’s multiple-comparisons test against the vehicle control. (G) CCK-8-based assessment of relative cell viability in CFPAC-1 cells treated with the indicated compounds for 24 or 48 h. Data are presented as mean ± SD from three independent biological experiments (n = 3). Statistical significance was assessed using two-way ANOVA followed by Tukey’s multiple-comparisons test.

Immunoblot analysis showed no substantial reduction in the abundance of G3BP1 and G3BP2, two key regulators of SG assembly, or TIA1, an SG-associated RBP, following treatment with the four compounds (Fig. 1E–F). These results indicate that the reduction in G3BP1-positive puncta was not accompanied by depletion of these major SG-associated proteins. We further assessed compound-associated changes in cellular metabolic activity and relative viability using the Cell Counting Kit-8 (CCK-8) assay. Analysis of the same dataset at 24 and 48 h revealed distinct effects of the compounds on cellular metabolic activity. Notably, HY-N0052A markedly reduced the background-corrected, normalized CCK-8 signal, approaching zero at both time points, whereas MB maintained substantially higher relative viability at concentrations effective in reducing SG accumulation (Fig. 1G). These findings raised concerns about the cytotoxic potential of HY-N0052A and supported the selection of MB for further investigation.

Given the genetic heterogeneity of PDAC, we next examined whether the effects of these compounds extended beyond CFPAC-1 cells by testing two additional PDAC cell lines, PANC-1 and MIA PaCa-2, which harbor distinct KRAS mutations (21). All four compounds reduced AS-induced G3BP1-positive puncta in both cell lines, with MB showing consistent activity (Fig. S2A–D). The abundance of these major SG-associated proteins remained largely unchanged in both PANC-1 and MIA PaCa-2 cells (Fig. S2E–H), indicating that the reduction in G3BP1-positive puncta was not accompanied by decreased expression of the examined SG components. Together, these findings identify MB as a reproducible modulator of stress-induced SG accumulation across multiple PDAC cell lines, with a more favorable cellular viability profile than HY-N0052A under the conditions examined.

### Methylene blue alters G3BP1 condensate formation and modulates G3BP1– Caprin1 association

Having identified MB as a reproducible suppressor of SG accumulation in PDAC cells, we next examined its effects on G3BP1 condensation, a key process underlying SG assembly. Because MB is a planar phenothiazine compound that can nonspecifically intercalate into nucleic acids, we first assessed whether MB directly affects the protein scaffold using a Ficoll 400-based reconstituted system devoid of RNA. Under these crowded cell-free conditions, purified polyhistidine (His)-tagged G3BP1 formed discrete, spherical condensates (Fig. 2A). Treatment with 10 μM MB markedly reduced the total number of condensates, while the few remaining assemblies displayed increased mean area and perimeter alongside decreased circularity (Fig. 2A–B) (22). This morphological divergence reflects an impairment of regular spherical droplet nucleation and an emergence of aberrant coalescent clusters. Critically, these data demonstrate that MB-mediated perturbation of G3BP1 phase separation operates intrinsically at the protein level, independent of RNA intercalation or non-specific nucleotide sequestration.

**Fig. 2.**
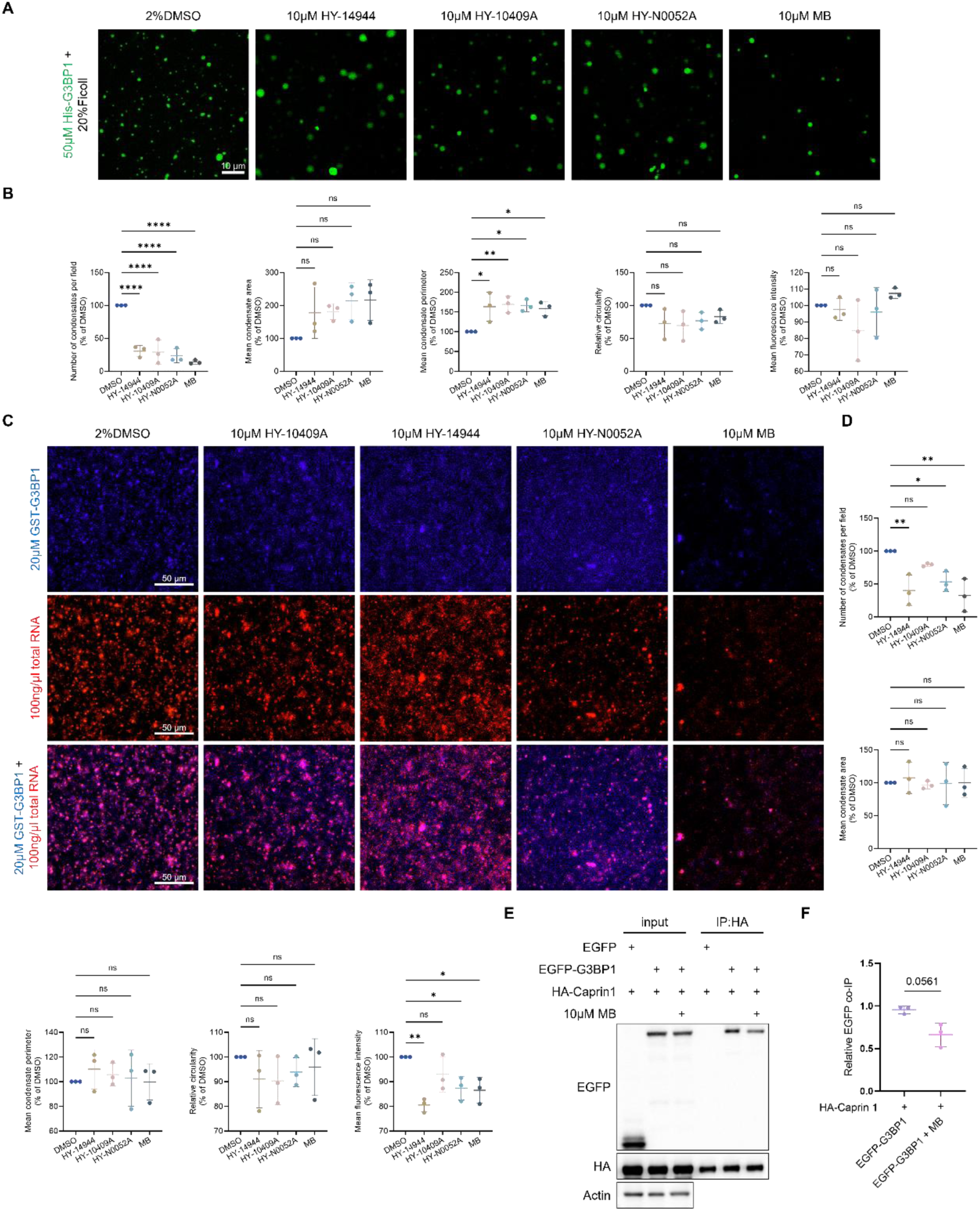
Methylene blue alters G3BP1 condensate formation and modulates G3BP1–Caprin1 association. (A) Representative confocal fluorescence images of His-G3BP1-WT condensates formed in a reconstituted system without added RNA containing 50 μM protein and 20% (w/v) Ficoll 400, in the presence of vehicle (2% DMSO) or the indicated compounds (10 μM). Scale bar, 10 μm. (B) Quantification of condensate number, mean area, perimeter, circularity, and mean fluorescence intensity in (A), normalized to the vehicle control (100%). (C) Representative fluorescence images of condensates formed by GST-G3BP1 (20 μM) and HEK-293T-derived total RNA (100 ng/μL) in the presence of vehicle (2% DMSO) or the indicated compounds (10 μM). GST-G3BP1 was labeled with Alexa Fluor 405, and RNA was stained with propidium iodide. Scale bar, 50 μm. (D) Quantification of condensate number, mean area, perimeter, circularity, and mean fluorescence intensity in (C), normalized to the vehicle control (100%). (E) Co-immunoprecipitation analysis of G3BP1–Caprin1 association in HEK-293T cells coexpressing HA-Caprin1 with EGFP vector or EGFP-G3BP1. Cells expressing EGFP-G3BP1 were treated with AS (1 mM) and vehicle or MB (10 μM) for 1 h before anti-HA immunoprecipitation and immunoblotting. (F) Quantification of co-immunoprecipitated EGFP-G3BP1 normalized to immunoprecipitated HA-Caprin1 and expressed relative to the AS-treated vehicle control. For (B) and (D), data are presented as mean ± SD from three independent LLPS reactions (n = 3). Statistical significance was determined using ordinary one-way ANOVA followed by Dunnett’s multiple-comparisons test against the corresponding DMSO control. For (F), data are presented as mean ± SD from three independent biological experiments (n = 3). Statistical significance was assessed using an unpaired two-tailed Welch’s t-test (P = 0.0561).

We next examined an RNA-containing reconstituted system using purified glutathione S-transferase (GST)-G3BP1 (20 μM) and total RNA isolated from HEK-293T cells (100 ng/μL). Under these conditions, MB reduced both the number and mean fluorescence intensity of G3BP1/RNA-positive condensates at 10 μM (Fig. 2C– D). To assess whether the GST tag itself contributed to condensate formation, we examined GST alone under identical conditions. GST did not form appreciable condensates in the presence of RNA, indicating that the condensation observed with GST-G3BP1 was not attributable to the GST tag alone (Fig. S3A). Increasing MB concentrations further reduced the abundance of detectable G3BP1/RNA-positive condensates (Fig. S3B–C). Together, these findings indicate that MB alters G3BP1 condensation in reconstituted systems both with and without added RNA.

We next tested whether MB alters the physical interaction between G3BP1 and Caprin1, which binds directly to the NTF2L hydrophobic groove. Co-immunoprecipitation in HEK-293T cells revealed that following AS exposure, MB co-treatment was accompanied by a modest, nonsignificant trend toward lower EGFP-G3BP1 retrieval with HA-Caprin1 *(P* = 0.0561; Fig. 2E–F). This subtle trend indicates that cellular-level SG suppression by MB cannot be attributed to a complete biochemical dissociation of Caprin1, but may instead arise from fine-tuning condensate nucleation thresholds and dynamic network remodeling.

Because MB possesses intrinsic redox activity and can modulate reactive oxygen species (ROS), we also examined whether its effects on SG accumulation extended to stress conditions other than AS exposure. Under nutrient deprivation or heat shock, MB reduced G3BP1-positive puncta in both CFPAC-1 and MIA PaCa-2 cells (Fig. S3D– G). These findings indicate that the suppression of SG accumulation by MB is not restricted to AS-induced oxidative stress. Collectively, our results show that MB alters G3BP1 condensation in reconstituted systems with and without added RNA, is associated with a nonsignificant trend toward reduced G3BP1 – Caprin1 co-immunoprecipitation, and reduces SG accumulation under multiple stress conditions.

### Molecular docking and dynamics simulations predict a potential interaction between methylene blue and the G3BP1 NTF2L domain

The observed effects of MB on G3BP1 condensation, together with the trend toward reduced G3BP1 – Caprin1 co-immunoprecipitation, prompted us to investigate its potential interaction with the NTF2L domain. Molecular docking was performed using Schrödinger Maestro and CB-Dock2. Both approaches placed MB within a similar hydrophobic groove of the NTF2L domain and identified partially overlapping sets of predicted interacting residues (Fig. 3A–B). To further evaluate this proposed interaction, we performed 100-ns molecular dynamics (MD) simulations of apo G3BP1 and the modeled G3BP1–MB complex.

**Fig. 3.**
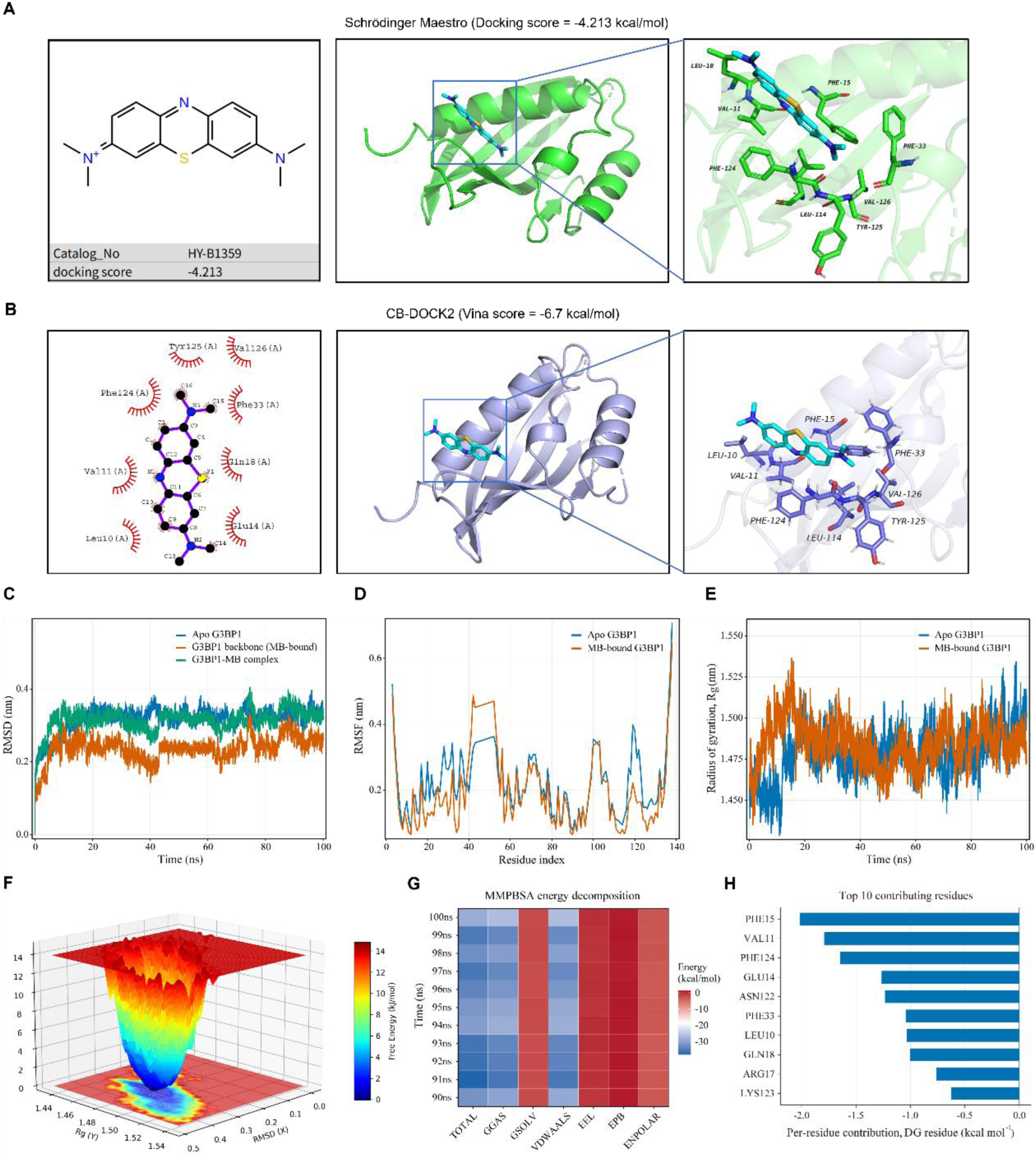
Molecular docking and dynamics simulations predict a potential interaction between methylene blue and the G3BP1 NTF2L domain. (A) Chemical structure of MB and its predicted binding pose within the G3BP1 NTF2L domain obtained using Schrödinger Maestro. The overall docking model and enlarged view of the predicted binding pocket are shown. Docking score, −4.213 kcal/mol. (B) Predicted MB-binding mode obtained using CB-Dock2, showing a two-dimensional ligand–residue interaction diagram, the overall docking pose, and an enlarged view of the binding pocket. Vina score, −6.7 kcal/mol. (C) RMSD profiles of apo G3BP1, the G3BP1 backbone in the MB-bound system, and the G3BP1–MB complex during 100-ns MD simulations. (D) Per-residue RMSF profiles of apo and MB-bound G3BP1. (E) Rg profiles of apo and MB-bound G3BP1 during the 100-ns MD simulations. (F) Gibbs free-energy landscape of the modeled G3BP1–MB complex as a function of RMSD and Rg. (G) Heatmap of MM/PBSA energy components calculated from the final 10 ns (90–100 ns) of the G3BP1–MB simulation, including total energy, gas-phase energy, solvation energy, van der Waals interactions, electrostatic interactions, polar solvation energy, and nonpolar solvation energy. (H) The ten residues with the most favorable contributions to the calculated G3BP1– MB interaction energy, identified by MM/PBSA per-residue energy decomposition.

In the simulation initiated from the docked structure, MB remained near the predicted NTF2L interaction surface throughout the sampled trajectory (Fig. S4A). Root-mean-square deviation (RMSD) analysis showed no sustained drift in the G3BP1 backbone during the later stages of the simulations (Fig. 3C). Root-mean-square fluctuation (RMSF) analysis revealed differences in local flexibility between apo and MB-associated G3BP1, with reduced fluctuations in regions encompassing residues 20– 40 and 112–130 and moderately increased fluctuations around residues 40–55 (Fig. 3D). These findings indicate differences in local conformational dynamics between the simulated systems.

Analysis of the radius of gyration (Rg) and solvent-accessible surface area (SASA) showed no major changes in overall protein compactness or solvent exposure during the simulations (Fig. 3E; Fig. S4B). Direction-resolved Rg analyses similarly showed no pronounced changes in global dimensions (Fig. S4C–D). Potential-energy, pressure, and temperature trajectories were monitored to assess simulation behavior (Fig. S4E– G). The free-energy landscape projected onto RMSD and Rg revealed a dominant conformational region sampled by the modeled G3BP1–MB complex (Fig. 3F; Fig. S4I).

We next evaluated the predicted interaction energetics using molecular mechanics/Poisson–Boltzmann surface area (MM/PBSA) analysis of the final 10 ns of the trajectory. The calculations yielded favorable interaction energies, with van der Waals interactions providing a major favorable contribution, whereas polar solvation opposed complex formation (Fig. 3G). Intermolecular hydrogen bonds between MB and G3BP1 were transient during the simulation (Fig. S4H). Per-residue energy decomposition identified Val11, Phe15, and Phe124 among the major favorable contributors to the modeled interaction (Fig. 3H; Fig. S4J).

Together, these computational analyses suggest that MB may interact with a hydrophobic groove within the G3BP1 NTF2L domain, with Val11, Phe15, and Phe124 emerging as potential contributors to the predicted interaction. The convergence of the docking predictions and MD-based energetic analyses provided a structural rationale for subsequent mutational studies of this region.

### Mutational analysis implicates the G3BP1 NTF2L hydrophobic region in the response to methylene blue

To investigate the functional relevance of the predicted NTF2L interaction region, we selected four hydrophobic residues, Val11, Phe15, Phe33, and Phe124, based on our computational analyses and previous structural and functional studies (23, 24). These residues contribute to a hydrophobic interaction pocket within the NTF2L domain involved in the recognition of G3BP1-binding partners, including Caprin1 and USP10. Whereas Val11, Phe15, and Phe124 were identified as favorable contributors in our MM/PBSA analysis, Phe33 was included based on its previously reported role in this interaction region. We therefore generated a quadruple alanine-substitution mutant (V11A/F15A/F33A/F124A; G3BP1-4A) to examine the contribution of this region to G3BP1 condensation and its response to MB (Fig. 4A). Recombinant His-tagged wild-type (WT) G3BP1 and G3BP1-4A were subsequently purified for biochemical analysis (Fig. S5A–B). Compared with WT G3BP1, the mutant showed lower protein recovery and additional coeluting protein bands during initial purification, consistent with previous reports of reduced recombinant protein yields for certain G3BP1 NTF2L mutants (24). To obtain a preparation suitable for subsequent condensation assays, the mutant-containing elution fractions were collected, concentrated, and subjected to molecular-weight-based filtration to reduce lower-molecular-weight contaminants.

**Fig. 4.**
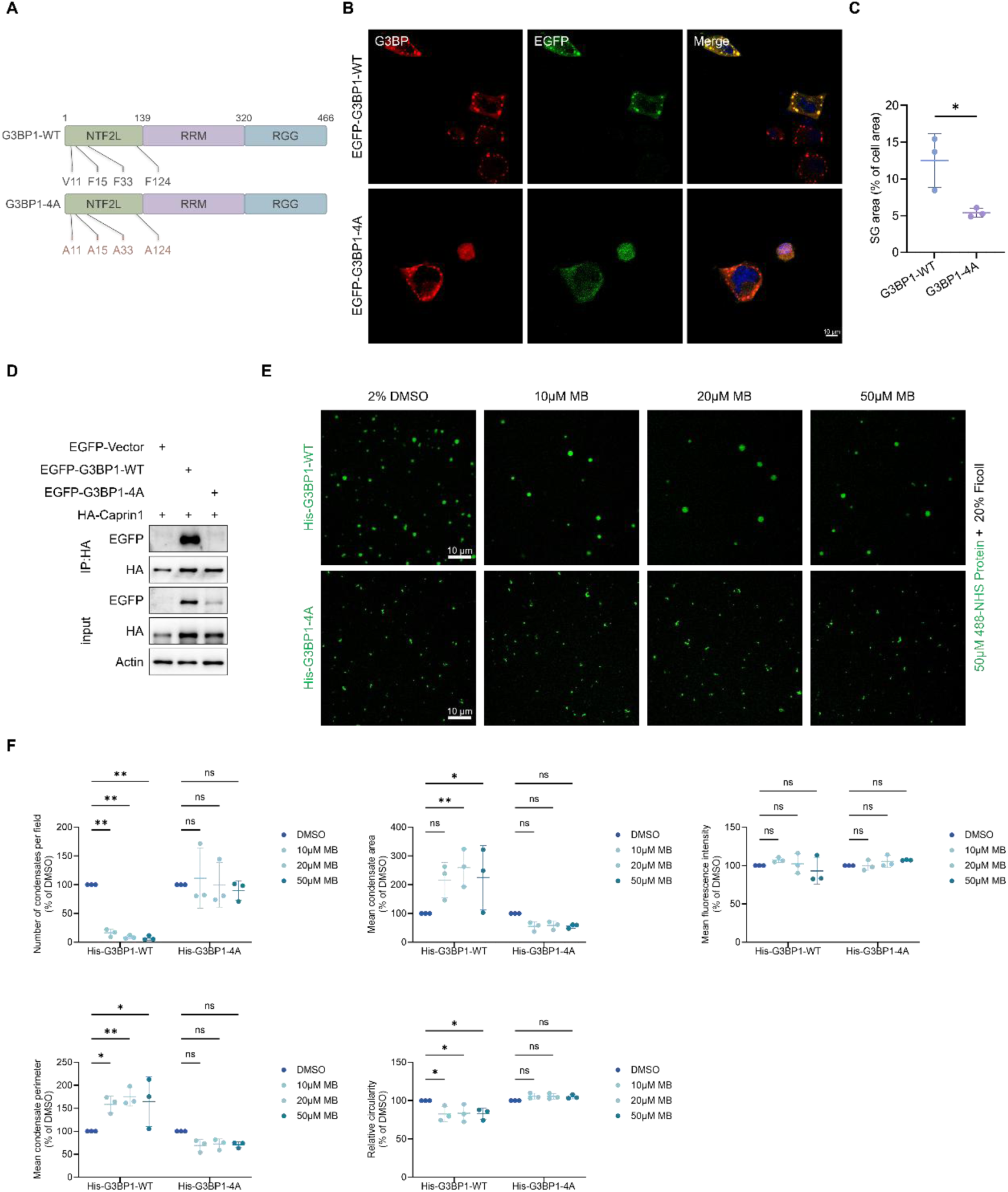
Mutational analysis implicates the G3BP1 NTF2L hydrophobic region in the response to methylene blue. (A) Schematic representation of G3BP1-WT and G3BP1-4A, highlighting the V11A, F15A, F33A, and F124A substitutions within the NTF2L domain. (B) Representative IF images of cells expressing EGFP-G3BP1-WT or EGFP-G3BP1-4A following AS (1 mM) treatment for 1 h. G3BP1 IF is shown in red, EGFP in green, and DAPI-stained nuclei in blue. Scale bar, 10 μm. (C) Quantification of SG accumulation in (B), expressed as the percentage of cellular area occupied by G3BP1-positive puncta. Data are presented as mean ± SD from three independent biological experiments (n = 3). Statistical significance was determined using an unpaired two-tailed Student’s t-test. (D) Co-immunoprecipitation analysis of G3BP1–Caprin1 association in HEK-293T cells coexpressing HA-Caprin1 with EGFP vector, EGFP-G3BP1-WT, or EGFP-G3BP1-4A. Cells were treated with AS (1 mM) for 1 h before anti-HA immunoprecipitation and immunoblotting. (E) Representative confocal fluorescence images of condensates formed by purified His-G3BP1-WT or His-G3BP1-4A (50 μM) in a reconstituted system without added RNA containing 20% (w/v) Ficoll 400, with vehicle (2% DMSO) or MB (10, 20, or 50 μM). Scale bar, 10 μm. (F) Quantification of condensate number, mean area, mean fluorescence intensity, perimeter, and circularity in (E). Values were normalized to the corresponding vehicle control for each protein (100%). Data are presented as mean ± SD from three independent LLPS reactions (n = 3). Statistical significance was determined using ordinary one-way ANOVA followed by Dunnett’s multiple-comparisons test against the corresponding vehicle control.

We first examined the effects of these substitutions on SG accumulation in cells. Following AS treatment, cells ectopically expressing G3BP1-4A still formed G3BP1-positive puncta, although their overall accumulation was reduced compared with cells expressing WT G3BP1 (Fig. 4B–C). We next assessed whether these substitutions affected the association between G3BP1 and Caprin1. Representative co-immunoprecipitation analysis showed substantially less detectable EGFP-G3BP1-4A associated with HA-Caprin1 than EGFP-G3BP1-WT under the conditions examined (Fig. 4D). These observations are consistent with previous studies implicating the NTF2L hydrophobic region in G3BP1–Caprin1 association and SG assembly.

We next compared the condensation properties of purified WT G3BP1 and G3BP1-4A under identical Ficoll 400-based molecular crowding conditions without added RNA. In the absence of MB, G3BP1-4A retained the ability to form detectable condensates. Although its condensate count was numerically lower than that of G3BP1-WT, none of the five measured basal condensation parameters differed significantly between the two proteins (Fig. S5C). We then examined the effects of increasing MB concentrations on both proteins under the same conditions. In the WT system, MB progressively reduced condensate number, accompanied by increases in the mean area and perimeter of the remaining condensates and a decrease in circularity (Fig. 4E–F). In contrast, G3BP1-4A showed no statistically significant changes in these parameters over the MB concentrations tested. Drug responses were evaluated relative to the corresponding vehicle control for each protein. These findings are consistent with a contribution of the NTF2L hydrophobic region to the response of G3BP1 condensates to MB, although a direct genotype-by-treatment interaction was not established by the present statistical analysis.

### Chronic oral administration of methylene blue is well tolerated under the tested regimens in wild-type mice

Having established that MB alters G3BP1 condensation and SG accumulation in reconstituted and cellular systems, we next evaluated its tolerability during prolonged administration in vivo. Previous studies have examined oral MB treatment in mice over extended periods (25, 26). To assess the regimens intended for subsequent studies of pancreatic disease, we administered MB through drinking water to WT C57BL/6J mice at a nominal dose of 10 mg/kg for 5 months (Fig. 5A). MB-treated mice showed no overt abnormalities in general appearance or gross pancreatic morphology compared with vehicle-treated controls (Fig. 5B). Body weight and the pancreas-to-body weight ratio were not significantly different between groups (Fig. 5C–D).

**Fig. 5.**
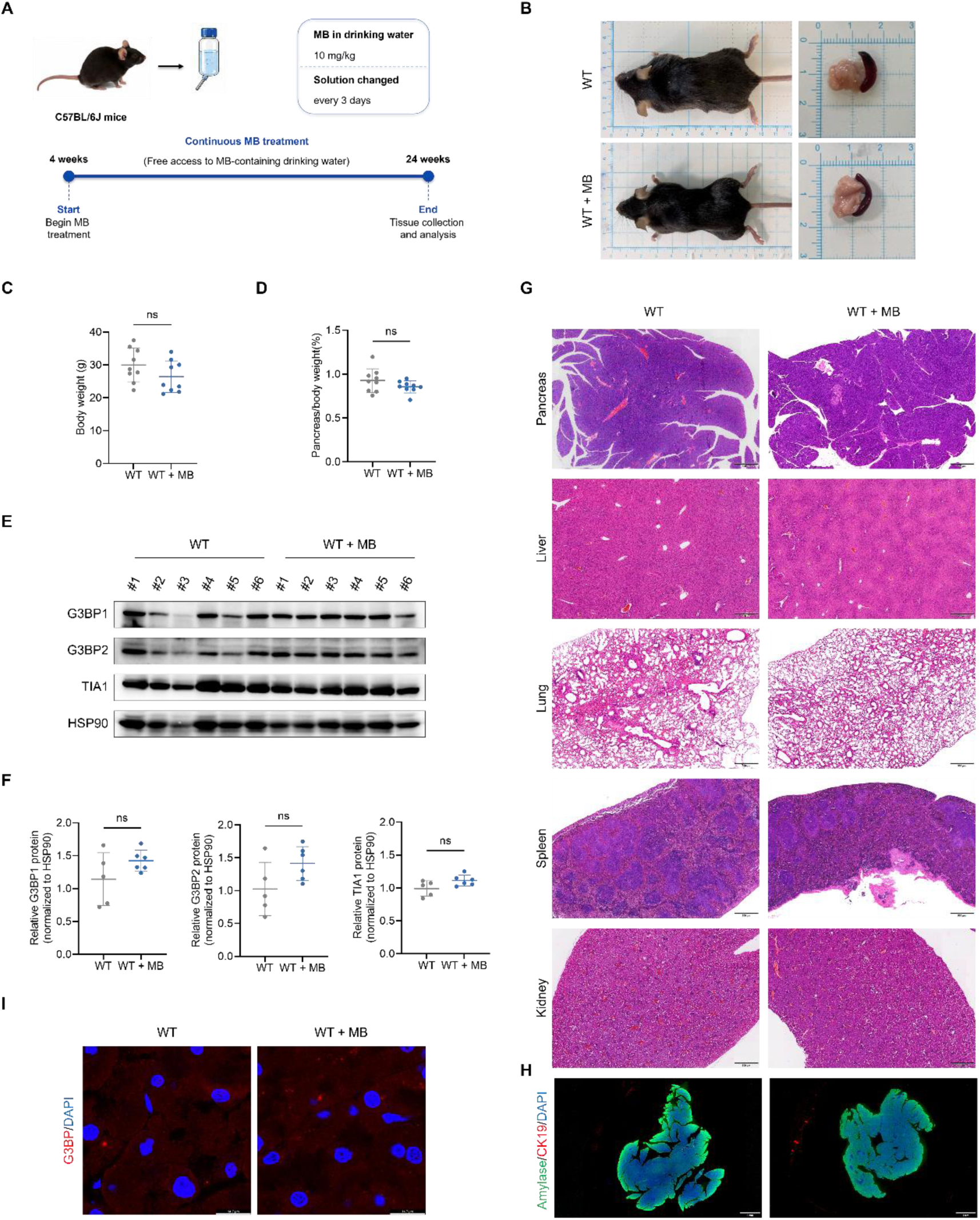
Chronic oral administration of methylene blue is well tolerated in wild-type mice under the tested regimen. (A) Schematic of the MB administration regimen. Four-week-old WT C57BL/6J mice received MB in drinking water at a nominal dose of 10 mg/kg for 5 months. Drinking water was replaced every 3 days. (B) Representative photographs of WT mice and excised pancreata from vehicle- and MB-treated groups. (C) Body weights of WT mice at the experimental endpoint. (D) Pancreas-to-body weight ratios of WT mice at the experimental endpoint. (E) Representative immunoblots of G3BP1, G3BP2, and TIA1 in pancreatic tissues from vehicle- and MB-treated WT mice. HSP90 served as the loading control. (F) Densitometric quantification of G3BP1, G3BP2, and TIA1 protein abundance in (E), normalized to HSP90. Six pancreatic protein samples per group were selected for immunoblot analysis; one WT control sample was excluded from quantification because of protein degradation (n = 5 WT control mice and n = 6 MB-treated mice). (G) Representative H&E-stained sections of the pancreas, liver, lung, spleen, and kidney from vehicle- and MB-treated WT mice. Scale bars, 500 μm. (H) Representative pancreatic tissue IF images showing amylase (green), CK19 (red), and DAPI (blue). Scale bars, 1 mm. (I) Representative pancreatic tissue IF images showing G3BP1 (red) and DAPI (blue). Scale bars, 14.7 μm. Data are presented as mean ± SD. For (C) and (D), n = 9 mice per group; for (F), n = 5 WT control mice and n = 6 MB-treated mice. Statistical significance was determined using an unpaired two-tailed Welch’s t-test. ns, not significant.

We next examined whether chronic MB administration affected the abundance of major SG-associated proteins or normal pancreatic tissue organization. Immunoblot analysis showed no significant changes in pancreatic G3BP1, G3BP2, or TIA1 abundance following MB treatment (Fig. 5E – F). Histological examination of the pancreas, liver, lung, spleen, and kidney revealed no obvious treatment-associated abnormalities in the tissue sections examined (Fig. 5G). IF staining for amylase and cytokeratin 19 (CK19) showed preservation of the normal acinar and ductal compartments, while G3BP1 staining remained predominantly diffuse without conspicuous accumulation of punctate structures in pancreatic tissue (Fig. 5H–I).

To further assess tolerability at a higher exposure level, we performed an additional study in WT mice receiving MB in drinking water at a nominal dose of 40 mg/kg for 3 months (Fig. S6A). Similar to the lower-dose regimen, no obvious differences in general appearance, gross pancreatic morphology, body weight, or pancreas-to-body weight ratio were observed relative to controls (Fig. S6B– D). Pancreatic G3BP1, G3BP2, and TIA1 abundance remained largely unchanged (Fig. S6E–F). Histological examination of major organs and IF analysis of pancreatic amylase, CK19, and G3BP1 revealed no overt treatment-associated alterations (Fig. S6G–I).

Together, these findings indicate that prolonged oral MB administration was generally well tolerated in WT mice under the regimens examined, without detectable changes in the measured body-weight, pancreatic protein-abundance, or tissue-morphology endpoints. These observations supported subsequent evaluation of MB in a genetically engineered mouse model of *Kras*-driven pancreatic disease.

### Methylene blue suppresses stress granule accumulation and attenuates *Kras*-driven pancreatic lesion progression in KC mice

Having established the effects of MB on G3BP1 condensation and SG assembly in vitro and in cultured cells, and confirmed its tolerability in WT mice under the tested regimens, we next investigated its effects on *Kras*-driven pancreatic disease in vivo. We employed *Pdx1-Cre; LSL-Kras^G12D^* (KC) mice, a well-established model that develops pancreatic intraepithelial neoplasias (PanINs) and recapitulates key features of early pancreatic tumorigenesis (27). Previous studies have implicated SG formation in *Kras*-driven pancreatic tumorigenesis and cellular stress adaptation (7, 8). KC mice received MB in drinking water at a nominal dose of 10 mg/kg for 5 months. MB treatment did not significantly affect body weight but reduced the pancreas-to-body weight ratio compared with untreated KC controls (Fig. 6A–C). Western blot analysis showed that the pancreatic protein levels of G3BP1, G3BP2, and TIA1 remained largely unchanged following MB administration (Fig. 6D–E).

**Fig. 6.**
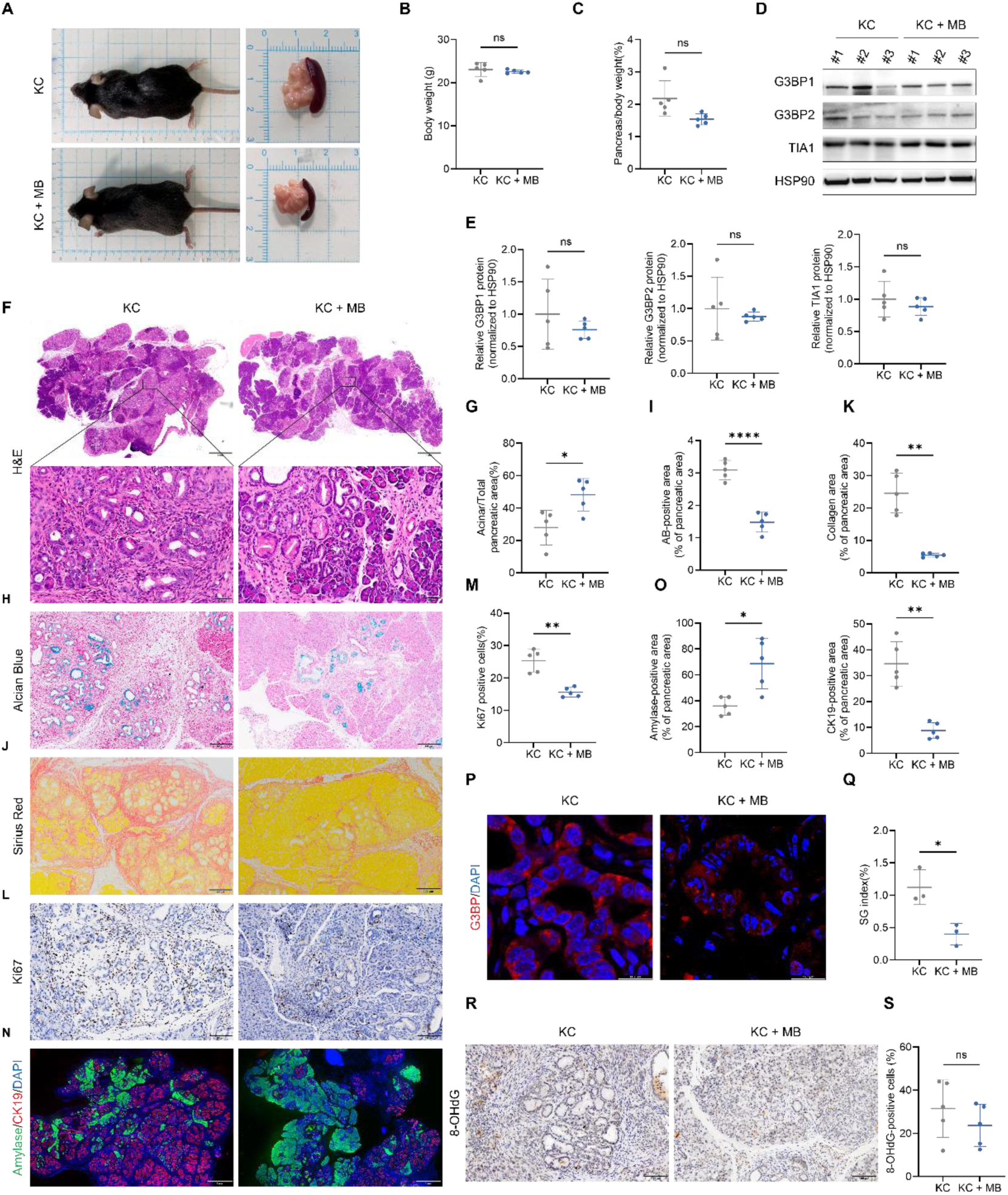
Methylene blue attenuates Kras-driven pancreatic preneoplastic lesions, stromal remodeling, and G3BP1-positive puncta in KC mice. (A) Representative photographs of KC mice and excised pancreata following administration of vehicle or MB in drinking water at a nominal dose of 10 mg/kg for 5 months. (B) Body weights of vehicle- and MB-treated KC mice at the experimental endpoint. (C) Pancreas-to-body weight ratios of vehicle- and MB-treated KC mice. (D) Representative immunoblots showing G3BP1, G3BP2, and TIA1 protein abundance in pancreatic tissues. HSP90 served as the loading control. (E) Densitometric quantification of the indicated proteins in (D), normalized to HSP90. (F) Representative H&E-stained pancreatic sections and enlarged views of the indicated regions. Scale bars, 1,000 μm (overview) and 50 μm (enlarged views). (G) Quantification of preserved acinar tissue area as a percentage of the total analyzed pancreatic tissue area. (H) Representative Alcian Blue-stained pancreatic sections showing mucin-positive ductal lesions. Scale bars, 200 μm. (I) Quantification of Alcian Blue-positive lesion area as a percentage of the total analyzed pancreatic tissue area. (J) Representative Sirius Red-stained pancreatic sections showing collagen deposition. Scale bars, 200 μm. (K) Quantification of the Sirius Red-stained area fraction, expressed as the percentage of red-stained area relative to the combined red- and yellow-stained areas. (L) Representative Ki-67 immunohistochemical staining of pancreatic tissues. Scale bars, 100 μm. (M) Quantification of Ki-67-positive cells as a percentage of the total evaluated cells. (N) Representative pancreatic tissue IF images showing amylase (green), CK19 (red), and DAPI (blue). Scale bars, 1000 μm. (O) Quantification of amylase-positive acinar area and CK19-positive ductal area, each expressed as a percentage of the analyzed pancreatic tissue area. (P) Representative confocal IF images showing G3BP1-positive puncta (red) and DAPI-stained nuclei (blue) in pancreatic tissues. Scale bars, 14.7 μm. (Q) Quantification of the pancreatic SG index, defined as the percentage of analyzed pancreatic tissue area occupied by G3BP1-positive puncta. (R) Representative immunohistochemical staining of 8-OHdG in pancreatic tissues. Scale bars, 100 μm. (S) Quantification of 8-OHdG-positive cells as a percentage of the total evaluated cells. Quantitative data are presented as mean ± SD (n = 5 mice per group), except for the SG index in (Q) (n = 3 mice per group). Histological area measurements were obtained from whole-section images, whereas three to five confocal fields were analyzed per mouse for tissue IF. Each mouse was considered an independent biological replicate. Statistical significance was determined using an unpaired two-tailed Welch’s t-test.

We next assessed the effects of MB on pancreatic lesion development and tissue remodeling. Hematoxylin and eosin (H&E) staining revealed substantial preservation of acinar tissue and reduced replacement by abnormal ductal structures in MB-treated KC mice, as confirmed by quantitative analysis of the preserved acinar area (Fig. 6F– G). Alcian Blue staining showed a significant reduction in mucin-positive ductal lesions (Fig. 6H–I), whereas Sirius Red staining demonstrated decreased collagen deposition (Fig. 6J–K). Consistently, Ki-67 immunohistochemistry revealed reduced proliferative activity in pancreatic tissues following MB treatment (Fig. 6L–M). IF analysis further showed increased amylase-positive acinar area and decreased CK19-positive ductal area (Fig. 6N–O). Together, these findings demonstrate that MB attenuates *Kras*-driven pancreatic lesion development, preserves pancreatic tissue architecture, and reduces stromal remodeling and cellular proliferation.

Consistent with its cellular activity, IF revealed abundant G3BP1-positive puncta throughout control KC pancreata, whereas their accumulation was significantly reduced in MB-treated KC mice, as quantified by the pancreatic SG index (Fig. 6P-Q). Importantly, this reduction occurred without altering the total abundance of G3BP1, G3BP2, or TIA1 (Fig. 6D-E). To evaluate whether tissue preservation was merely a secondary consequence of systemic ROS scavenging, we quantified 8-hydroxy-2’-deoxyguanosine (8-OHdG) (28). 8-OHdG levels showed no significant differences between groups (Fig. 6R-S), indicating that reduced lesion burden and SG puncta occurred in the absence of measurable changes in this classic oxidative DNA damage biomarker. While these findings demonstrate that chronic MB treatment simultaneously attenuates pancreatic SG burden and early histopathology in vivo, they reflect a robust phenotypic correlation rather than an isolated cause-and-effect proof.

To further validate these findings, we examined an independent cohort of KC mice receiving MB at a nominal dose of 40 mg/kg for 3 months. Consistent with the lower-dose regimen, MB treatment reduced the pancreas-to-body weight ratio without significantly affecting body weight, while pancreatic G3BP1, G3BP2, and TIA1 protein levels remained unchanged (Fig. S7A–E). Histological analyses again showed greater preservation of acinar tissue, reduced mucin-positive lesion area and collagen deposition, and decreased Ki-67 staining (Fig. S7F–M). Amylase-positive area was increased, while CK19-positive area showed a reduction that did not reach statistical significance (Fig. S7N–O). Furthermore, MB markedly reduced pancreatic G3BP1-positive puncta and the corresponding SG index (Fig. S7P–Q), whereas no statistically significant difference in the percentage of 8-OHdG-positive cells was observed between vehicle- and MB-treated mice (Fig. S7R–S).

As summarized in Fig. 7, MB is proposed to modulate the hydrophobic interaction region of the G3BP1 NTF2L domain, potentially influencing G3BP1 – Caprin1 association and altering G3BP1 condensation without substantially reducing the abundance of major SG-associated proteins. These effects are consistent with the reduced SG accumulation observed in PDAC cells and pancreatic tissues, accompanied by attenuation of *Kras*-driven pancreatic lesion development, cellular proliferation, and stromal remodeling. Together, our findings highlight G3BP1-associated condensate assembly as a pharmacologically accessible process and provide a framework for further investigation of MB-mediated modulation of SG dynamics in pancreatic disease.

**Fig. 7.**
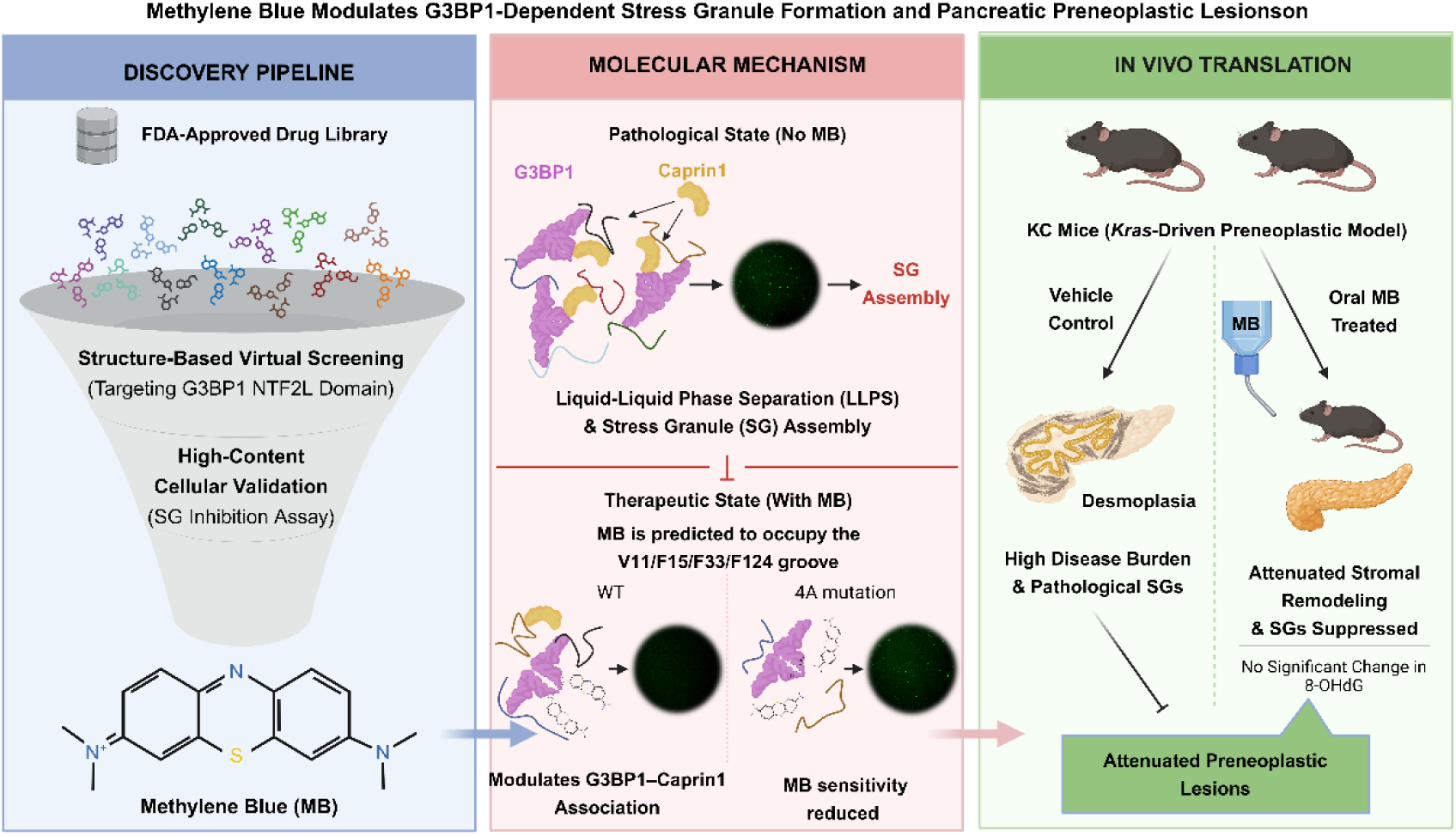
Proposed model of methylene blue-mediated regulation of G3BP1 condensation and attenuation of *Kras*-driven pancreatic lesions. Schematic summary of the discovery strategy, proposed molecular mechanism, and in vivo effects of MB. Structure-based virtual screening of an FDA-approved drug library targeting the G3BP1 NTF2L domain, followed by cellular validation, identified MB as a modulator of SG accumulation. Molecular docking and mutational analyses implicate a hydrophobic region containing V11, F15, F33, and F124 in the response of G3BP1 to MB. MB alters G3BP1 condensate formation and reduces cellular SG accumulation, whereas the G3BP1-4A mutant exhibits reduced sensitivity to MB. These effects are accompanied by a trend toward decreased G3BP1 – Caprin1 association. In *Kras*-driven KC mice, chronic MB administration attenuates pancreatic preneoplastic lesions, preserves acinar tissue, reduces stromal remodeling, and decreases pancreatic G3BP1-positive puncta. The model illustrates a potential mechanism linking MB-mediated modulation of G3BP1 condensation to the attenuation of pancreatic preneoplastic pathology.

## Discussion

SGs are increasingly recognized as important regulators of cellular stress adaptation and potential therapeutic targets in cancer. In PDAC, oncogenic KRAS signaling and persistent environmental stress create conditions in which SG-associated pathways may support tumor cell survival and disease progression (7, 8). Despite growing interest in pharmacologically targeting SG assembly, effective chemical modulators remain limited (19). Drug repurposing offers an attractive strategy for addressing this gap (29). Here, we identified MB as a modulator of G3BP1-associated condensation through an integrated approach combining structure-guided drug screening, cellular assays, biochemical reconstitution, computational modeling, mutational analysis, and animal studies. MB suppressed SG accumulation across multiple PDAC cell lines and distinct stress conditions and altered G3BP1 condensation in reconstituted systems with or without added RNA. Co-immunoprecipitation analysis further revealed a nonsignificant trend toward reduced G3BP1–Caprin1 association. These effects were accompanied by reduced SG accumulation and attenuation of pancreatic lesions in *Kras*-driven KC mice. Together, our findings support G3BP1-associated condensate assembly as a pharmacologically accessible process with relevance to pancreatic tumorigenesis.

G3BP1 and G3BP2 are central regulators of SG assembly, coordinating multivalent protein–RNA interactions that promote condensation under stress (12–14). The N-terminal NTF2L domain contributes to G3BP dimerization and serves as an interaction platform for regulatory proteins, including Caprin1 and USP10 (12, 16, 18). Previous studies have established the functional importance of its hydrophobic interaction region (23, 24). Our molecular docking and dynamics simulations localized a plausible MB interaction pose within the conserved hydrophobic groove formed by Val11, Phe15, Phe33, and Phe124. In support of this predicted binding site, the G3BP1-4A mutant exhibited reduced Caprin1 association and diminished SG formation in cells, while preserving basal condensation competence in vitro without significant baseline deviations from wild-type G3BP1. Crucially, whereas MB exerted pronounced morphological effects on WT G3BP1 condensates, G3BP1-4A showed reduced sensitivity to MB across identical concentrations. While these mutational findings functionally implicate the NTF2L hydrophobic cleft in mediating MB sensitivity, we emphasize that they do not exclude possible allosteric influences or off-target cellular interactions, which remain to be resolved by structural co-crystallography.

The complementary biochemical systems provide further insight into the effects of MB on condensate assembly. RNA is a major regulator of G3BP1 condensation and SG organization (13, 14, 30). Although MB reduced G3BP1/RNA-positive condensates in the RNA-containing system, it also altered condensation of purified G3BP1 under Ficoll 400-based molecular crowding conditions without added RNA. Thus, its activity cannot be explained solely by interference with externally supplied RNA. Notably, fewer condensates were accompanied by increased mean area and perimeter and decreased circularity of the remaining condensates in the Ficoll 400 system, indicating changes in condensate organization rather than a uniform reduction in all morphological parameters. Together with the observed trend toward reduced G3BP1– Caprin1 association, these findings support a model in which MB modifies G3BP1 condensation by influencing its molecular interaction network.

MB has well-documented redox activity and can influence mitochondrial function and cellular signaling (31, 32). Its effects on purified G3BP1 condensation and SG accumulation under multiple stress conditions suggest that its activity is not restricted to AS-induced oxidative stress, although redox-dependent contributions cannot be excluded. Interestingly, MB has been reported to accelerate the liquid-to-gel transition of tau condensates (33), highlighting the context-dependent effects of MB on biomolecular condensation.

In KC mice, MB administration attenuated pancreatic preneoplastic lesions, preserved acinar architecture, and reduced collagen deposition and cellular proliferation. These changes were accompanied by fewer pancreatic G3BP1-positive puncta without substantial changes in G3BP1, G3BP2, or TIA1 abundance. Similar findings were observed under an independent administration regimen. No significant difference in 8-OHdG-positive cell percentages was detected, indicating that the histological improvements were not accompanied by a measurable reduction in this oxidative DNA damage marker (28). In WT mice, MB was generally well tolerated based on the evaluated endpoints, without overt histological abnormalities in the examined organs. Blue discoloration of selected tissues and urine was observed, consistent with previously reported effects of MB administration and excretion (25, 34–37). Nevertheless, additional pharmacokinetic and safety studies are required to characterize its long-term systemic effects.

Several mechanistic complexities and experimental limitations must be explicitly considered. First, methylene blue is a polypharmacological molecule with well-documented redox activity, mitochondrial actions, and nitric oxide synthase inhibition (31, 38–40). Although the comparable 8-OHdG levels observed between vehicle and MB cohorts indicate that pancreatic lesion suppression is not simply a byproduct of decreased steady-state oxidative DNA damage, we cannot rule out that MB-driven systemic metabolic or stromal modulations contribute to disease attenuation. Second, in preneoplastic KC pancreata, lesion progression, chronic tissue inflammation, and SG assembly form an interconnected vicious cycle (7, 41). Disentangling whether MB-mediated reduction in G3BP1 puncta is a primary driving trigger of delayed PanIN progression or a secondary hallmark of healthier, acinar-preserved tissue remains challenging in pharmacological studies; definitive causal dissection will require tissue-specific G3BP1 phase-separation-deficient knock-in models. Third, while the in silico docking, MD trajectories, and G3BP1-4A mutational analyses consistently converge on the NTF2L hydrophobic cleft, solution biophysical measurements (e.g., ITC or SPR) and high-resolution co-crystal structures will be indispensable to definitively quantify the thermodynamic binding affinity and stereochemical coordinates. Additional limitations of the animal experiments include unequal sex distributions between treatment groups in the 5-month cohorts, the lack of drinking-water consumption measurements, and the absence of blinded image analysis. Potential sex-related confounding therefore cannot be excluded. Furthermore, the two MB administration regimens differed in both nominal dose and treatment duration and were not designed to establish a dose – response relationship. Despite these limitations, our findings provide a foundation for investigating G3BP1-associated condensation as a pharmacologically modifiable process in early *Kras*-driven pancreatic disease (2, 24, 42).

## Materials and methods

### Schrödinger Maestro Molecular Docking

The human G3BP1 structure (PDB ID: 8V1L) was retrieved from the Protein Data Bank and prepared using the Protein Preparation Wizard in Schrödinger Maestro 12.8. Hydrogen atoms were added, crystallographic water molecules and accessory chains (chains B–F) were removed to isolate the functional Chain A monomer, and the receptor was energy-minimized using the OPLS2005 force field with an RMSD threshold of 0.30 Å. A receptor grid measuring 20 × 20 × 20 Å was generated using the Receptor Grid Generation module and centered on the canonical hydrophobic peptide-binding pocket of the NTF2L monomer occupied by Y9M.

The 2,783 compounds from the FDA-approved drug library were prepared using LigPrep, including hydrogen addition, three-dimensional structure generation, and energy minimization. Structure-based virtual screening was performed using the Glide Virtual Screening Workflow in extra-precision (XP) mode. Of the 2,783 compounds, 2,738 yielded valid docking results, whereas 45 did not produce valid results because docking was unsuccessful. Compounds with valid results were ranked by docking score, and the predicted binding poses of selected candidates were examined for interactions within the G3BP1 NTF2L domain. The complete available docking results are deposited in Zenodo, and the identities of the 45 unsuccessful compounds are provided in Supplementary Table S1.

### CB-Dock2 Molecular Docking

Potential binding pockets on the G3BP1 surface were identified using the CurPocket algorithm implemented in CB-Dock2 (43, 44). MB was docked into the identified pockets using AutoDock Vina (45), and the resulting binding poses were evaluated based on their docking scores and predicted molecular interactions. The predicted interactions were visualized in two dimensions using LigPlot+ (46) and in three dimensions using PyMOL.

### Molecular Dynamics Simulations and Binding Energy Calculations

All-atom MD simulations of apo G3BP1 and the modeled G3BP1–MB complex were performed using GROMACS v2025.04. The protein was described using the CHARMM36m force field, and MB parameters were generated using Sobtop with the General AMBER Force Field (GAFF). Each system was solvated with TIP3P water, maintaining a minimum distance of 1.0 nm between the solute and the simulation box boundaries, and neutralized with Na⁺ or Cl⁻ ions. Following steepest-descent energy minimization, the systems were equilibrated for 2 ns under constant number, volume, and temperature (NVT) conditions and 2 ns under constant number, pressure, and temperature (NPT) conditions. Production simulations were then conducted for 100 ns under periodic boundary conditions at 300 K and 1 bar, using a 2-fs integration time step. Temperature and pressure were controlled using V-rescale and Berendsen coupling, respectively. Short-range electrostatic and van der Waals interactions were truncated at 1.0 nm, long-range electrostatic interactions were treated using the particle mesh Ewald method, and bonds involving hydrogen atoms were constrained using LINCS.

Trajectories were analyzed using GROMACS tools to assess RMSD, RMSF, Rg, SASA, intermolecular hydrogen bonds, potential energy, temperature, and pressure. Hydrogen bonds were defined by a donor–acceptor distance cutoff of 3.5 Å and a donor–hydrogen–acceptor angle greater than 120°. Gibbs free-energy landscapes were constructed using RMSD and Rg as reaction coordinates. Binding energies were estimated using gmx_MMPBSA with MMPBSA.py v1.6.4, based on trajectory frames from the final 10 ns (90–100 ns) of the production simulation. The calculated binding energies were decomposed into van der Waals, electrostatic, polar solvation, and nonpolar solvation contributions, followed by per-residue energy decomposition.

### Mice

C57BL/6J mice were purchased from Jiangsu GemPharmatech Co., Ltd. *Kras*-driven KC (*LSL-Kras^G12D/+^; Pdx1^Cre/+^*) mice were generated by crossing *LSL-Kras^G12D/+^* mice with *Pdx1^Cre/+^*mice. All mice were maintained under specific pathogen-free (SPF) conditions. All animal experiments were conducted in accordance with applicable national guidelines and approved by the Animal Ethics Committee of Xuzhou Medical University (approval no. 202411T009).

### Cell lines and Culture Conditions

The human PDAC cell lines CFPAC-1, PANC-1, and MIA PaCa-2 were obtained from the Cell Resource Center, Institute of Basic Medical Sciences. HEK-293T cells were provided by Prof. Yong Liu (Xuzhou Medical University). CFPAC-1 cells were maintained in RPMI 1640 medium (Gibco, Cat. no. C11875500BT) supplemented with 10% fetal bovine serum (FBS; PN-Biotech GmbH, Cat. no. ST30-3302) and 1% penicillin–streptomycin (VICMED, Cat. no. VC2003). PANC-1, MIA PaCa-2, and HEK-293T cells were cultured in DMEM (Gibco, Cat. no. C11995500BT) supplemented with 10% FBS and 1% penicillin–streptomycin. All cells were maintained at 37°C in a humidified incubator containing 5% CO₂.

### Stress Induction and Drug Treatments

For compound screening, CFPAC-1 cells were seeded in 96-well plates or 6-well plates containing glass coverslips 48 h before treatment. CFPAC-1, PANC-1, and MIA PaCa-2 cells were subsequently subjected to the indicated stress conditions.

For oxidative stress induction, AS (NaAsO₂; Sigma, Cat. no. S7400) was dissolved in PBS to prepare a 100 mM stock solution. When cells reached approximately 80% confluence, AS was diluted in the corresponding culture medium to a final concentration of 1 mM. Candidate compounds (10 μM) or vehicle were added simultaneously with AS, and cells were incubated at 37°C for 1 h. Unstressed controls were included in parallel. Following treatment, cells were washed with PBS and processed for IF staining.

For heat-shock experiments, CFPAC-1 and MIA PaCa-2 cells were exposed to 44°C for 1 h in the presence of 10 μM MB or an equivalent volume of vehicle. Cells were subsequently washed with PBS and processed for IF staining.

For nutrient-deprivation experiments, CFPAC-1 and MIA PaCa-2 cells were cultured in medium lacking glucose, glutamine, HEPES, and sodium pyruvate (Servicebio, Cat. no. G4528) for 5 and 16 h, respectively. During the final hour of nutrient deprivation, cells were treated with 10 μM MB or vehicle. Following treatment, cells were washed with PBS and processed for IF staining.

### Immunofluorescence Staining of Cultured Cells

Following the indicated treatments, cells were washed with PBS and fixed with 4% paraformaldehyde (PFA; VICMED, Cat. no. VIH100) for 15 min at room temperature. Cells were permeabilized with 0.2% Triton X-100 (Sigma, Cat. no. T8787) for 10 min at 4°C and blocked with 3% bovine serum albumin (BSA; VICMED, Cat. no. VIC018) for 1 h at room temperature. Cells were then incubated with an anti-G3BP1 antibody (Abcam, Cat. no. ab56574; 1:200 in 3% BSA) overnight at 4°C.

After three washes with PBS for 10 min each, cells were incubated with Alexa Fluor 594-conjugated goat anti-mouse IgG (H+L) (Invitrogen, Cat. no. A-11005; 1:200) for 1 h at room temperature. Nuclei were subsequently counterstained with DAPI (Solarbio, Cat. no. C0065) for 15 min without an intervening wash. Following three additional washes with PBS for 10 min each, coverslips were mounted using antifade mounting medium (Solarbio, Cat. no. S2100).

### Quantification of Cellular SGs

G3BP1-positive cytoplasmic puncta were quantified from IF images using Fiji/ImageJ with a customized macro-assisted workflow (47). Individual cells were manually delineated as regions of interest (ROIs), and G3BP1-positive puncta were identified using a semi-manual thresholding and particle-analysis procedure. Identical analysis settings were applied across treatment groups within each experiment.

The SG index was calculated for each cell as follows:

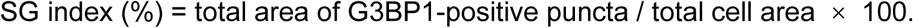

SG index values from individual cells were averaged to obtain one value per image. Image-level values were subsequently averaged within each independent biological experiment to generate one value per treatment condition. Cells and images from the same experiment were treated as nested measurements rather than independent biological replicates. The number of independent experiments, data presentation, and statistical comparisons are specified in the corresponding figure legends.

### Western Blot Analysis

Cells and tissues were lysed in RIPA buffer containing 150 mM NaCl, 50 mM Tris-HCl (pH 7.4), 0.1% SDS, 1 mM EDTA, 1% NP-40, and 1% sodium deoxycholate, supplemented with protease and phosphatase inhibitors. Lysates were clarified by centrifugation, and protein concentrations were determined using a BCA protein assay. Equal amounts of protein (20 μg per lane) were separated by SDS-PAGE and transferred onto polyvinylidene fluoride (PVDF) membranes.

Membranes were blocked with 5% nonfat milk in TBST for 1 h at room temperature and incubated with primary antibodies overnight at 4 ° C. The primary antibodies included G3BP1 (Proteintech, Cat. no. 13057-2-AP; 1:1000), G3BP1 (Abcam, Cat. no. ab56574; 1:3000), G3BP2 (Abcam, Cat. no. ab86135; 1:5000), TIA1 (Proteintech, Cat. no. 12133-2-AP; 1:2000), EIF2S1 (Abcam, Cat. no. ab32157; 1:5000), phospho-EIF2S1 (Abcam, Cat. no. ab5369; 1:5000), HSP90 (Proteintech, Cat. no. 13171-1-AP; 1:20000), β-actin (Proteintech, Cat. no. 66009-1-Ig; 1:20000), HA tag (Proteintech, Cat. no. 51064-2-AP; 1:2000), and GFP (Invitrogen, Cat. no. A-11122; 1:2000). After three washes with TBST for 5 min each, membranes were incubated with the appropriate HRP-conjugated secondary antibodies (Proteintech, Cat. nos. SA00001-1 and SA00001-2; 1:5000) for 1 h at room temperature.

Protein bands were visualized using NcmECL SuperUltra chemiluminescent substrate (NCM Biotech, Cat. no. P10100) and a Tanon-5200 imaging system. Band intensities were quantified using Fiji/ImageJ and normalized to HSP90. The number of independent biological replicates and the statistical analyses are specified in the corresponding figure legends.

### CCK-8 Cell Metabolic Activity and Relative Viability Assays

Cellular metabolic activity and relative viability were assessed using the Cell Counting Kit-8 (CCK-8; VICMED, Cat. no. VC5001L). CFPAC-1 cells were seeded in 96-well plates at a density of 4,000 cells per well and cultured for 24 h before treatment with homoharringtonine (HY-14944), fedratinib hydrochloride hydrate (HY-10409A), sanguinarine chloride (HY-N0052A), or methylene blue (MB; HY-B1359) at the indicated concentrations. Vehicle-treated cells served as controls.

CCK-8 measurements were performed after 24 and 48 h of compound treatment. At each time point, 10 μL of CCK-8 reagent was added per well, followed by incubation at 37°C for 1 h. Absorbance was measured at 450 nm using a Cytation 3 multimode reader (BioTek Instruments). Compound-matched cell-free controls were included for background correction, and the corresponding blank absorbance was subtracted from each measurement.

Separate plates were prepared during the same cell-seeding session for measurements at 24 and 48 h after compound treatment. Thus, each time point was assessed using a distinct set of wells rather than repeated measurements of the same wells. Three independent biological experiments were performed (n = 3), with six technical replicate wells per condition in each experiment. Technical replicate measurements were averaged to generate one value per treatment condition at each time point within each independent experiment. The resulting biological replicate values were used for statistical analysis.

### Construction of G3BP1-4A Mutants

The G3BP1 quadruple mutant (G3BP1-4A), containing the V11A, F15A, F33A, and F124A substitutions, was generated using a homologous recombination-based cloning strategy. DNA fragments containing the designed mutations and overlapping homologous sequences were generated by PCR and assembled into the corresponding expression vectors using homologous recombination reagents (Beyotime Biotechnology, China). Recombinant plasmids were subjected to DNA sequencing by GENEWIZ (China), which confirmed the intended substitutions. Sequence-verified constructs were used for subsequent cellular and recombinant protein experiments.

### Plasmids and Transfection

Cells were seeded in 10-cm dishes at 60–70% confluence. For transfection, 6 μg of plasmid DNA and 6 μL of Lipo8000 transfection reagent (Beyotime, Cat. no. C0533) were diluted in 1 mL of Opti-MEM (Gibco, Cat. no. 31985070) and mixed gently. After incubation at room temperature for 20 min, the transfection mixture was added dropwise to the cells. The medium was replaced with fresh complete medium after 6– 8 h. Subsequent experiments were performed at the indicated times after transfection.

The plasmids used in this study are listed below.

### Co-Immunoprecipitation

HEK-293T cells were co-transfected with HA-Caprin1 and EGFP vector, EGFP-G3BP1-WT, or EGFP-G3BP1-4A plasmids. For the MB treatment experiment (Fig. 2E– F), cells expressing EGFP-G3BP1-WT and HA-Caprin1 were treated with AS (1 mM) together with vehicle or MB (10 μM) for 1 h. For the WT/4A comparison (Fig. 4D), cells expressing the corresponding G3BP1 constructs and HA-Caprin1 were exposed to AS (1 mM) for 1 h without MB treatment. Cells were harvested 72 h after transfection and lysed in NETN buffer (50 mM Tris-HCl, 150 mM NaCl, 1 mM EDTA, and 1% NP-40) supplemented with protease and phosphatase inhibitors. Lysates were incubated with gentle rotation at 4°C for 2 h and clarified by centrifugation at 20,000 × g for 20 min at 4°C.

For each sample, 40 μL of clarified lysate was mixed with 10 μL of 5× SDS loading buffer and retained as the input fraction. The remaining lysate was incubated with anti-HA magnetic beads (Thermo Fisher Scientific, Cat. no. 88836) overnight at 4°C with gentle rotation. The beads were washed sequentially with TBST, NETN buffer, and NETN buffer containing 2% Tween-20, with each washing step performed twice. Bound proteins were eluted in 1 × SDS loading buffer at 95 ° C for 5 min. Input and immunoprecipitated fractions were analyzed by Western blotting using antibodies against HA and GFP.

Co-immunoprecipitated EGFP-G3BP1 was normalized to immunoprecipitated HA-Caprin1. Relative G3BP1 – Caprin1 association was expressed against the corresponding vehicle-treated WT control for Fig. 2F and the AS-treated WT reference for Fig. 4D.

### Protein Expression and Purification

For GST-G3BP1 purification, the pGEX-4T-1-G3BP1-GST construct was transformed into T7 Express competent *Escherichia coli* (Weidibio, Cat. no. EC2080). Bacterial cultures were grown to an OD_600_ of approximately 0.8 and induced with 0.6 mM isopropyl β-D-1-thiogalactopyranoside (IPTG; Beyotime, Cat. no. T1416-5G) at 16°C for 20 h. Cells were lysed using a bacterial protein extraction reagent (ACE, Cat. no. BR0005-02), and the clarified lysates were incubated with GST affinity resin (Beyotime, Cat. no. P2262) overnight at 4 ° C. The resin was washed according to the manufacturer’s instructions, and GST-G3BP1 was eluted with buffer containing 50 mM Tris-HCl, 150 mM NaCl, and 10 mM reduced glutathione (pH 8.0). The eluted protein was concentrated and buffer-exchanged into sodium bicarbonate buffer (pH 8.3) using an Amicon Ultra centrifugal filter with a 50-kDa molecular-weight cutoff (MWCO; Merck Millipore) for subsequent fluorescent labeling.

For His-tagged protein purification, the pET28a-G3BP1-His construct encoding G3BP1-WT and a corresponding sequence-verified pET28a-based construct encoding G3BP1-4A were independently transformed into Rosetta *E. coli*. Bacterial cultures were grown to an OD_600_ of approximately 0.6 and induced with 0.6 mM IPTG at 16°C for 20 h. Cells were lysed using the bacterial protein extraction reagent (ACE, Cat. no. BR0005-02), and the clarified lysates were incubated with His-tag affinity resin (Beyotime, Cat. no. P2226) overnight at 4°C. The purification buffers contained 50 mM sodium phosphate and 300 mM NaCl (pH 8.0), supplemented with the indicated concentrations of imidazole. His-G3BP1-WT was sequentially washed with buffers containing 5 mM and 8 mM imidazole and eluted with buffers containing 25 mM, 50 mM, and 75 mM imidazole. In contrast, His-G3BP1-4A was washed with 5 mM imidazole and eluted with 50 mM imidazole.

Protein-containing elution fractions were collected and subjected to centrifugal ultrafiltration using Amicon Ultra centrifugal filters (Merck Millipore) at 4°C and 4,000 rpm for 30 min. Membranes with MWCOs of 30 and 50 kDa were used for His-G3BP1-WT and His-G3BP1-4A, respectively. The proteins were concentrated and buffer-exchanged into sodium bicarbonate buffer (pH 8.3) for subsequent fluorescent labeling. Protein purification profiles were evaluated by SDS-PAGE.

### In Vitro Condensation (LLPS) Assays

For RNA-containing condensation assays, purified GST and GST-G3BP1 were fluorescently labeled with Alexa Fluor 405 NHS ester (Invitrogen, Cat. no. A30000) in sodium bicarbonate buffer (pH 8.3) for 1 h at room temperature in the dark. The labeling reaction was quenched with 200 mM Tris buffer (pH 8.0). Total RNA was extracted from HEK-293T cells using TRIzol reagent (TransGen Biotech, Cat. no. ET111-01-V2) and stained with propidium iodide (PI; MedChemExpress, Cat. no. HY-D0815).

RNA-containing condensation reactions were assembled in low-binding tubes at a final volume of 10 μL, containing purified GST-G3BP1 (20 μM) and total RNA (100 ng/μL) in 1× TBS buffer (20 mM Tris-HCl, 150 mM NaCl, pH 7.6). GST alone was included as a control. Candidate compounds or vehicle were added at the indicated concentrations. After gentle mixing, the samples were transferred to chambers formed between glass slides and coverslips separated by double-sided spacers (Sigma, Cat. no. GBL654002). Condensates were imaged using a Zeiss Axio Observer 7 fluorescence microscope with a 20× objective.

For condensation assays without added RNA, purified His-G3BP1-WT and His-G3BP1-4A were fluorescently labeled with Alexa Fluor 488 NHS ester (Invitrogen, Cat. no. A20000) in sodium bicarbonate buffer (pH 8.3) for 1 h at room temperature in the dark. Labeling was quenched with 200 mM Tris buffer (pH 8.0). A 50% (w/v) Ficoll 400 stock solution was prepared from Ficoll 400 powder (Sigma, Cat. no. F2637-10G) and diluted to a final concentration of 20% (w/v) in the condensation reactions. Reactions contained 50 μM protein in 1× TBS buffer (20 mM Tris-HCl, 150 mM NaCl, pH 7.6), together with the indicated compounds or vehicle. Samples were mixed by gentle pipetting at room temperature and imaged using an LSM 880 confocal laser scanning microscope (Carl Zeiss).

The concentrations of G3BP1 and RNA used in the RNA-containing condensation assays were selected based on published studies and preliminary experiments evaluating different combinations of protein and RNA concentrations. Under the conditions tested, higher RNA concentrations resulted in dense, precipitate-like material that was unsuitable for reliable quantification of discrete condensates, whereas lower RNA concentrations yielded too few detectable condensates to distinguish compound-associated effects. Accordingly, 20 μM GST-G3BP1 and 100 ng/μL total RNA were selected to provide quantifiable condensate formation and facilitate comparisons among treatment conditions.

Condensate number, mean area, perimeter, circularity, and mean fluorescence intensity were quantified using Fiji/ImageJ. Three to five fields of view were analyzed per independent condensation reaction, and field-level measurements were averaged to obtain one value per reaction. Fields acquired from the same reaction were not considered independent replicates. The numbers of independent experiments, normalization procedures, and statistical analyses are specified in the corresponding figure legends.

### Mice Treatments

MB (Sigma, Cat. no. 457250) was dissolved in 1× PBS to prepare a 2 mg/mL stock solution. After complete dissolution, the solution was filtered, aliquoted, and stored at −80°C until use. Four-week-old WT C57BL/6J and KC mice were used for the indicated experiments. Mice were randomly allocated to vehicle- and MB-treated groups within the respective experimental cohorts.

For the 5-month regimen, the WT vehicle group included 2 females and 7 males, whereas the WT MB group included 6 females and 3 males. The KC vehicle group included 4 females and 1 male, whereas the KC MB group included 5 females. For the 3-month regimen, both WT groups included 4 females and 2 males per group, and both KC groups included 4 females and 3 males per group. No formal a priori sample-size calculation was performed. Group sizes were determined based on previous experimental experience, animal availability, and the feasibility of the planned analyses.

For oral administration, the MB stock solution was diluted in sterile drinking water to achieve the nominal dosing regimens of 10 mg/kg for 5 months or 40 mg/kg for 3 months. Control mice received drinking water without MB. MB-containing drinking water was protected from light and freshly prepared every 3 days. Body weights were recorded weekly throughout the treatment period. Drinking-water consumption was not monitored; therefore, the stated MB doses represent nominal target doses rather than directly measured daily intake. The 10 mg/kg and 40 mg/kg regimens were evaluated in independent cohorts with different treatment durations and were not designed as a formal dose–response comparison.

At the experimental endpoint, mice were euthanized, and pancreata were photographed and weighed. Pancreas-to-body weight ratios were calculated using the corresponding terminal body weights. Pancreatic tissues and other indicated organs were collected for protein analysis, histological examination, immunohistochemistry, and IF. Individual mice were considered independent biological replicates, with group sizes reported for each endpoint in the corresponding figure legends.

### Immunohistochemistry

Immunohistochemical staining of pancreatic tissues was performed by Servicebio (Wuhan, China). Paraffin-embedded pancreatic sections (3.5 μm) were deparaffinized in xylene and rehydrated through a graded ethanol series. Heat-induced antigen retrieval was performed using citrate buffer (pH 6.0) for Ki-67 staining or Tris-EDTA buffer (pH 9.0) for 8-OHdG staining. Endogenous peroxidase activity was quenched with 3% hydrogen peroxide for 15 min at room temperature, followed by blocking to reduce nonspecific antibody binding.

Sections were incubated overnight at 4 ° C with anti-Ki-67 (Servicebio, Cat. no. GB111141; rabbit; 1:1000) or anti-8-OHdG (Servicebio, Cat. no. GB150115; mouse; 1:200) primary antibodies. Following PBS washes, Ki-67 staining was detected using an HRP-conjugated goat anti-rabbit IgG (H+L) secondary antibody, whereas 8-OHdG staining was detected using an HRP-conjugated goat anti-mouse IgG (H+L) secondary detection system. Immunoreactivity was visualized using 3,3′-diaminobenzidine (DAB), followed by hematoxylin counterstaining, differentiation, bluing, dehydration, clearing, and mounting.

Whole pancreatic tissue sections were scanned for quantitative analysis. Ki-67-positive nuclei were quantified using a custom Fiji/ImageJ macro (v61) incorporating hematoxylin–DAB color separation, nuclear segmentation, and DAB intensity-based classification. The nuclear segmentation threshold was set to 55, with a particle area range of 25–3000 pixels². DAB intensity thresholds of 12 and 65 were used to classify weakly and strongly stained nuclei, respectively.

8-OHdG-positive nuclei were quantified using a custom automated image analysis pipeline. Hematoxylin and DAB signals were separated by Ruifrok–Johnston color deconvolution. Nuclei were segmented using a hematoxylin optical density threshold of 0.18, an area range of 10 – 2000 pixels ² , and marker-controlled watershed segmentation. For each image, the DAB positivity threshold was defined as max(1.8 × p50, 0.12), where p50 represents the median DAB optical density within the valid tissue region. Nuclei were considered positive when at least 20% of nuclear pixels exceeded this threshold. Identical analysis rules were applied across experimental groups.

Positive nuclei were expressed as percentages of the total evaluated nuclei. Each mouse contributed one quantitative value and was treated as an independent biological replicate. Sample sizes and statistical analyses are provided in the corresponding figure legends.

### Histological Staining

Pancreatic tissues and other indicated organs were fixed in 4% paraformaldehyde (VICMED, Cat. no. VIH100) for 24 h, dehydrated, and embedded in paraffin. Pancreatic tissues were sectioned at 3.5 μm, whereas other organs were sectioned at 5 μm. H&E, Alcian Blue, and Sirius Red staining were performed by Servicebio (Wuhan, China) according to its established staining procedures.

For histological staining, paraffin sections were deparaffinized in two changes of xylene for 20 min each, followed by rehydration through two changes of absolute ethanol for 5 min each and 75% ethanol for 5 min, and rinsing with water.

For H&E staining, sections were stained with hematoxylin (Servicebio H&E staining kit, Cat. no. G1003) for 3 – 5 min, differentiated, blued, and rinsed. Sections were subsequently passed through 85% and 95% ethanol for 5 min each, stained with eosin for 5 min, dehydrated in three changes of absolute ethanol for 5 min each, cleared in two changes of xylene for 5 min each, and mounted.

For Alcian Blue staining, sections were stained with Alcian Blue staining solution A (Servicebio, Cat. no. G1027) for 10–15 min, rinsed with water, and counterstained with nuclear staining solution B for 3 min. Following rinsing, sections were dehydrated in three changes of absolute ethanol for 5 min each, cleared in two changes of xylene for 5 min each, and mounted.

For Sirius Red staining, sections were immersed in Sirius Red staining solution (Servicebio, Cat. no. G1018) for 8 min, rapidly dehydrated through two or three changes of absolute ethanol, cleared in xylene for 5 min, and mounted with neutral mounting medium.

### Quantitative Evaluation of Pancreatic Lesions

Whole pancreatic sections stained with H&E, Alcian Blue, or Sirius Red were scanned and analyzed using Fiji/ImageJ or custom Python scripts.

In H&E-stained sections, morphologically preserved acinar regions were manually delineated, and their areas were expressed as percentages of the total pancreatic tissue area. For Alcian Blue staining, mucin-positive regions were identified using color-based segmentation with the following criteria: B − R ≥ 20, G − R ≥ 0, V = 45– 252, and S ≥ 25. Alcian Blue-positive area was calculated relative to the total tissue area.

Sirius Red-stained sections were analyzed using HSV-based color classification. Red-stained regions were identified using hue values ≤ 28 ° or ≥ 335 °, a saturation threshold of 75, and a value threshold of 40. The red-stained area fraction was calculated as the red-stained area divided by the sum of red- and yellow-stained areas.

Measurements were obtained from whole-section images and summarized to obtain one value per mouse. Individual mice were treated as independent biological replicates. Sample sizes and statistical analyses are specified in the corresponding figure legends.

### Tissue Immunofluorescence

Paraffin-embedded pancreatic sections (3.5 μm) were deparaffinized in xylene and rehydrated through a graded ethanol series. Heat-induced antigen retrieval was performed in sodium citrate buffer (pH 6.0). Sections were permeabilized with 0.2% Triton X-100 for 10 min at 4°C and blocked with 3% BSA for 1 h at room temperature. Sections were incubated with the indicated primary antibodies overnight at 4°C. After washing with PBS, the corresponding fluorescence-conjugated secondary antibodies were applied for 1 h at room temperature. Nuclei were counterstained with DAPI for 15 min at room temperature. Following PBS washes, sections were mounted using antifade mounting medium.

Primary antibodies included anti-G3BP1 (Abcam, Cat. no. ab56574; 1:200), anti-cytokeratin 19 (CK19; Abcam, Cat. no. ab52625; 1:200), and anti-amylase (Santa Cruz Biotechnology, Cat. no. sc-46657; 1:200). Secondary antibodies included Alexa Fluor 594-conjugated goat anti-mouse IgG (H+L) (Invitrogen, Cat. no. A-11005; 1:200), Alexa Fluor 488-conjugated goat anti-mouse IgG (H+L) (Invitrogen, Cat. no. A-11001; 1:200), and Alexa Fluor 594-conjugated goat anti-rabbit IgG (H+L) (Invitrogen, Cat. no. A-11012; 1:200), as appropriate.

Fluorescence images were acquired using a Leica TCS SP5 confocal laser scanning microscope (Leica Microsystems). Three to five fields of view were analyzed per mouse using Fiji/ImageJ. Amylase-positive acinar areas and CK19-positive ductal areas were expressed as percentages of the total pancreatic tissue area analyzed within the acquired fields.

Pancreatic G3BP1-positive puncta were quantified from tissue IF images using Fiji/ImageJ. Fluorescence thresholding was used to identify discrete G3BP1-positive punctate signals while excluding background and diffuse fluorescence. The pancreatic SG index was defined as the total area occupied by G3BP1-positive puncta divided by the analyzed pancreatic tissue area, multiplied by 100.

Three to five confocal fields were examined per mouse, and field-level measurements were summarized to obtain one value per animal. Each mouse was treated as an independent biological replicate, whereas fields from the same mouse were considered within-animal measurements. Group sizes and statistical analyses are provided in the corresponding figure legends.

### Statistical Analysis

Quantitative data are presented as mean ± SD unless otherwise indicated. Statistical analyses were performed using GraphPad Prism. The number of independent biological replicates (n), statistical tests, and multiple-comparison procedures, where applicable, are specified in the corresponding figure legends.

For cell-based assays, independent experimental runs were considered biological replicates, whereas individual cells, microscopy images, and technical replicate wells within the same experiment were treated as nested or technical measurements. For in vitro condensation assays, independent reactions were considered experimental replicates, and measurements from multiple fields were summarized at the reaction level before statistical analysis. For animal studies, each mouse was considered an independent biological replicate; tissue sections and microscopy fields from the same mouse were not treated as independent replicates. For pancreatic histology and tissue IF, image acquisition and quantitative analyses using Fiji/ImageJ or custom Python scripts were performed without blinding to treatment-group allocation.

All statistical tests were performed as indicated in the figure legends. A two-sided P value < 0.05 was considered statistically significant. Statistical significance is denoted as *P < 0.05, **P < 0.01, ***P < 0.001, and ****P < 0.0001; ns, not significant.

## Acknowledgments

This work was supported by the National Natural Science Foundation of China (Grant Nos. 32600850, 82203311, 82573779), the Major Project of Basic Science (Natural Science) Research in Colleges and Universities of Jiangsu Province (Grant No. 24KJD320006 and 25KJB180016), the Basic Research Program of Jiangsu (Grant No. BK20220670), the Jiangsu Province Capability Improvement Project through Science, Technology and Education (NO. CXZX202234), the Open Project of Key Laboratory of Science and Engineering for the Multi-modal Prevention and Control of Major Chronic Diseases, Ministry of Industry and Information Technology (Grant No. MCD-2023-1-01), the Research Foundation of Xuzhou Medical University (Grant Nos. D2023004, D2023012, D2022034, D2025006), the Science and Technology project of Xuzhou (Grant No. KC23003), the Fundamental Research Funds for the Central Universities (No. 2024QN11068), ‘ Double Innovation ’ Doctor (No. 140924007), ‘ Double Innovation ’ talents projects of Jiangsu Province, the China Postdoctoral Science Foundation (GZB20240550, 2024M752406), Shanghai Postdoctoral Excellence Program (2023597), the Jiangsu Provincial Innovation and Entrepreneurship Team (Education Category), the Major Project of Basic Science (Natural Science) Research in Jiangsu Higher Education Institutions (Category A) (Grant No. 25KJA180006) and the Jiangsu Province Postgraduate Research and Practice Innovation Program (Grant Nos. KYCX25_3315 and 26CXJH5214). The funders had no roles in study design, data collection and analysis, decision to publish, or preparation of the manuscript.

## Author contributions

Conceptualization was the responsibility of WJ.G, ZJ.Z, J.X and WH.Z. Methodology was the responsibility of ZJ.Z, J.X, WH.Z, QD.L, LL.C, XY.W, YF.Z, X.L, Y.H, N.W, LY.M, JT.X, RH.Y, JK.D, C.H, HX.Z, YR.Z, CL.Y, MY.L, X.S, QL.Z, YR.C, SM.Z, XH.H. The investigation was conducted by WJ.G, ZJ.Z, J.X. The original draft was written by ZJ.Z and J.X. The reviewing and editing were performed by WJ.G, TT.W, and C.W. Fund acquisition was carried out by WJ.G, X.L, YF.Z. Supervision was the responsibility of WJ.G.

## Data and Code Availability

Data supporting this study are provided in the article and its supporting information. Unprocessed images and numerical source data are available from the corresponding author upon reasonable request.

Computational files, including docking preparation files, docking configurations, predicted poses and scores, and molecular dynamics input, parameter, and analysis files, are available in Zenodo (https://doi.org/10.5281/zenodo.23229272). Of the 2,783 compounds included in the initial virtual screening, 2,738 yielded valid docking results, which are available in the repository. The remaining 45 compounds did not produce valid docking results, and their identities are listed in Supplementary Table S1.

## Declaration of generative AI and AI-assisted technologies in the writing process

During the preparation of this manuscript, the authors used ChatGPT (OpenAI) to improve the clarity and language of existing draft text. All AI-assisted text was reviewed and edited by the authors, who take full responsibility for the final content.

## Supplementary figure legends

**Fig. S1.**
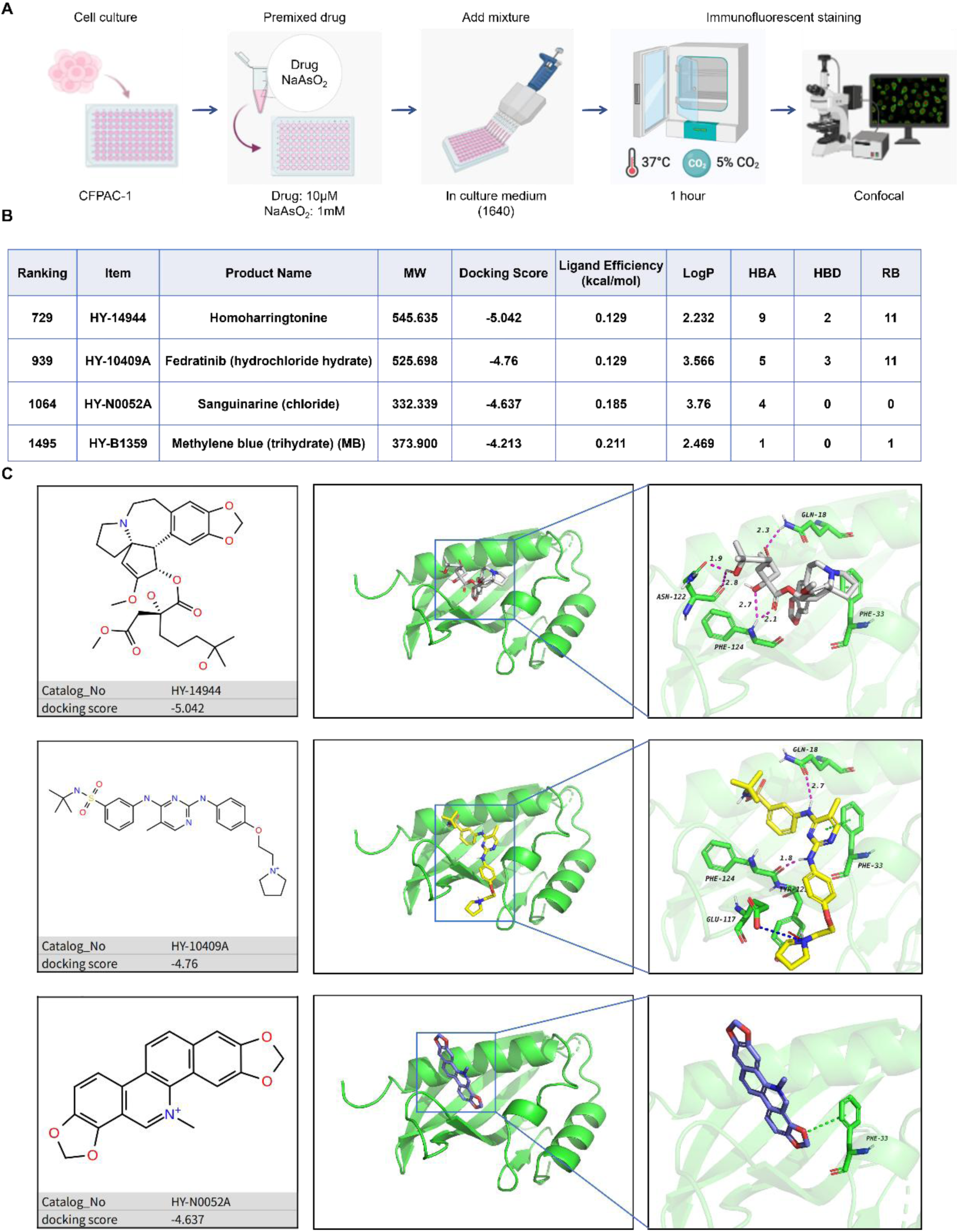
Docking-guided screening and predicted interactions of candidate compounds with the G3BP1 NTF2L domain. (A) Schematic overview of the cell-based screening workflow. CFPAC-1 cells were treated with the indicated compounds (10 μM) and AS (1 mM) for 1 h, followed by IF staining and confocal imaging of G3BP1-positive puncta. (B) Virtual screening ranks, docking scores, and molecular properties of four selected compounds—homoharringtonine (HY-14944), fedratinib hydrochloride hydrate (HY-10409A), sanguinarine chloride (HY-N0052A), and MB (HY-B1359)—obtained using Schrödinger Glide. (C) Chemical structures and predicted binding poses of HY-14944, HY-10409A, and HY-N0052A within the G3BP1 NTF2L domain. Overall docking models and enlarged views of the predicted binding pockets are shown. The corresponding docking scores are −5.042, −4.760, and −4.637 kcal/mol, respectively.

**Fig. S2.**
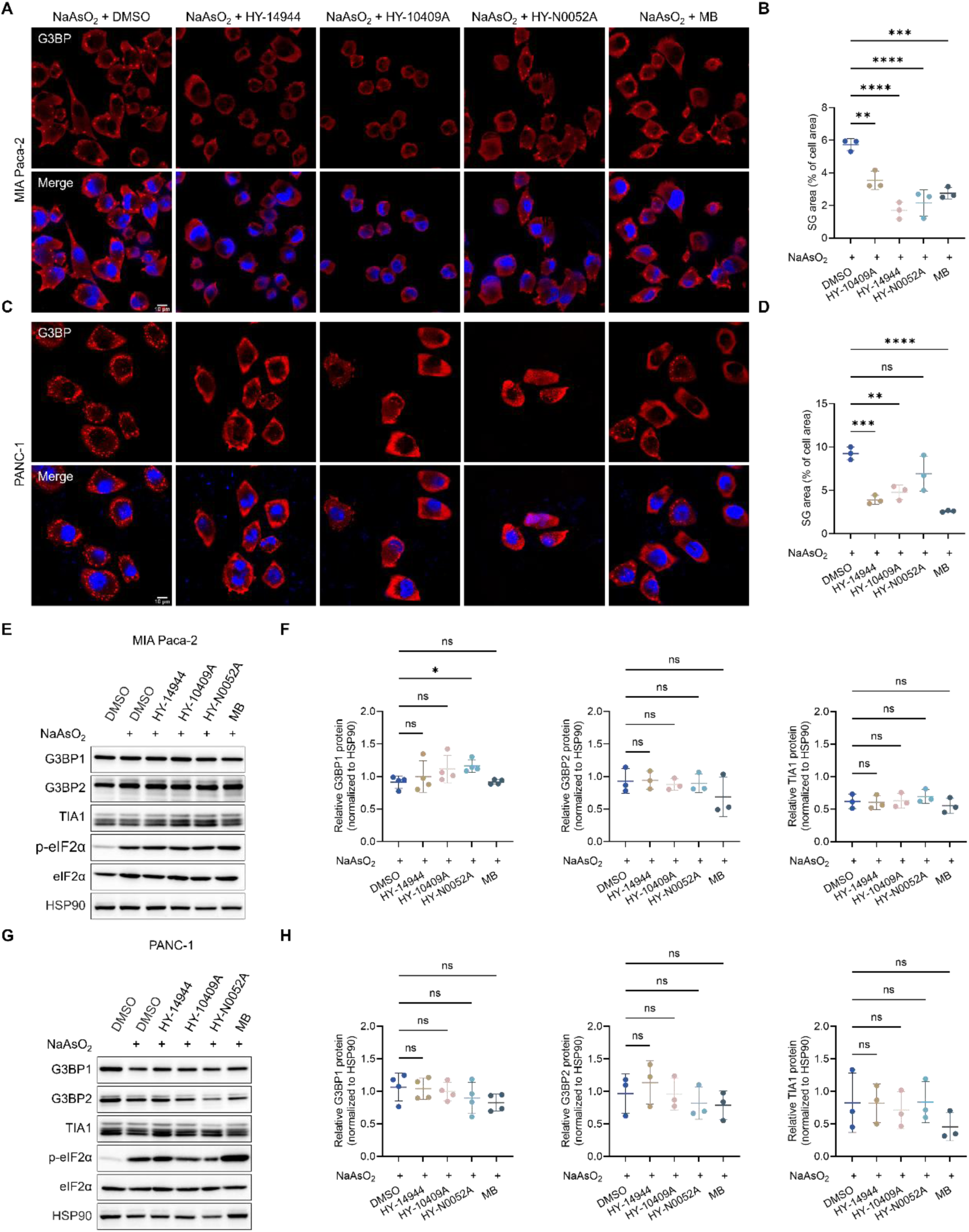
Validation of the effects of candidate compounds on stress granule accumulation in additional PDAC cell lines. (A) Representative IF images of G3BP1-positive SGs in MIA PaCa-2 cells treated with AS (1 mM) and vehicle or the indicated compounds (10 μM) for 1 h. G3BP1 is shown in red and DAPI-stained nuclei in blue. Scale bar, 10 μm. (B) Quantification of SG accumulation in (A), expressed as the percentage of cellular area occupied by G3BP1-positive puncta. (C) Representative IF images of G3BP1-positive SGs in PANC-1 cells treated under the same conditions as in (A). Scale bar, 10 μm. (D) Quantification of SG accumulation in (C), using the SG index described in (B). (E) Representative immunoblots of G3BP1, G3BP2, TIA1, phosphorylated eIF2α (p-eIF2α), and total eIF2α in MIA PaCa-2 cells following the indicated treatments. HSP90 served as the loading control. (F) Densitometric quantification of G3BP1, G3BP2, and TIA1 protein abundance in (E), normalized to HSP90. (G) Representative immunoblots of G3BP1, G3BP2, TIA1, p-eIF2α, and total eIF2α in PANC-1 cells following the indicated treatments. HSP90 served as the loading control. (H) Densitometric quantification of G3BP1, G3BP2, and TIA1 protein abundance in (G), normalized to HSP90. Data are presented as mean ± SD. For SG quantification in (B) and (D), n = 3 independent biological experiments. For immunoblot quantification in (F) and (H), G3BP1 was analyzed in four independent biological experiments (n = 4), whereas G3BP2 and TIA1 were analyzed in three independent biological experiments (n = 3). Statistical significance was determined using ordinary one-way ANOVA followed by Dunnett’s multiple-comparisons test against the corresponding AS-treated vehicle control.

**Fig. S3.**
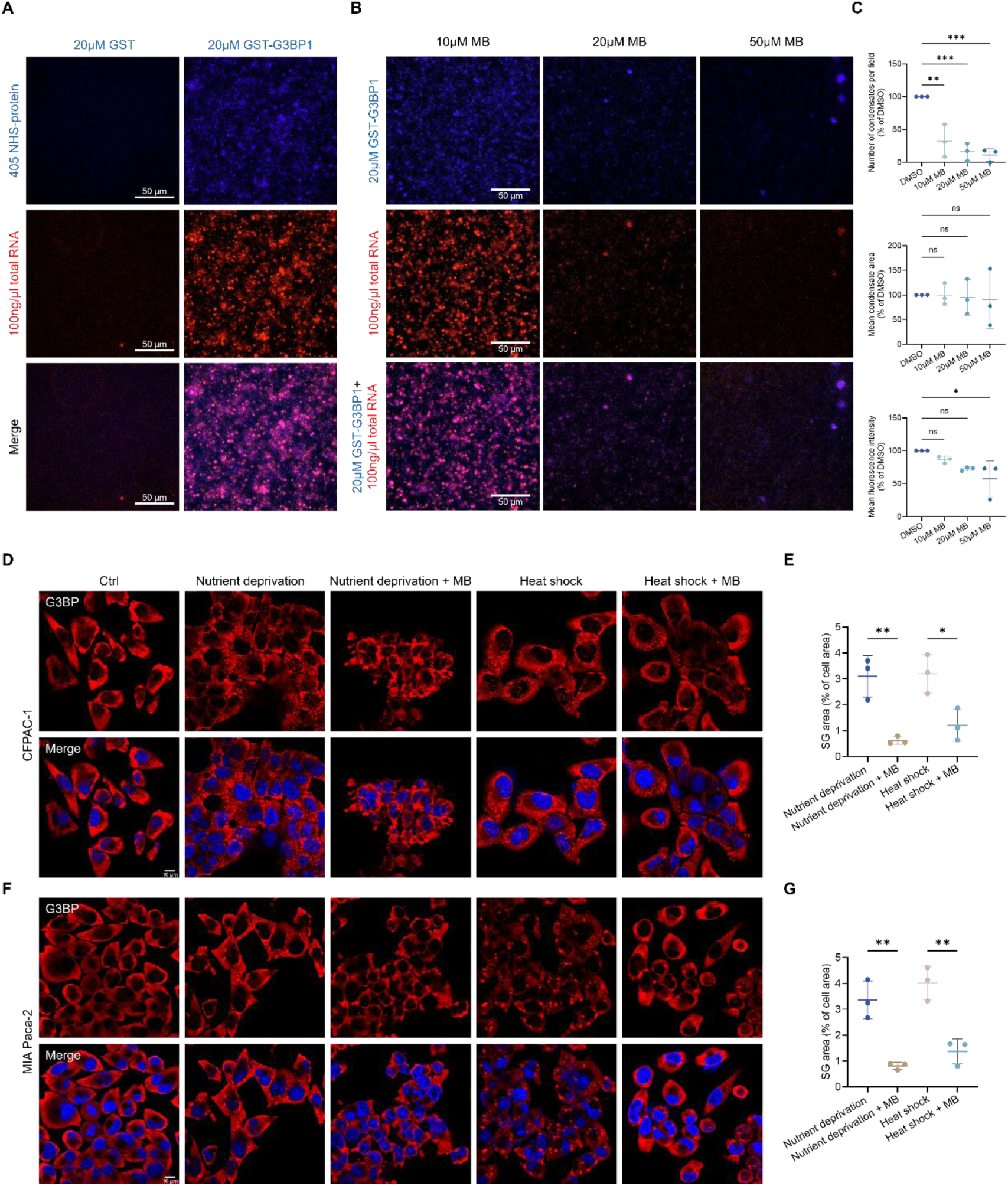
Methylene blue reduces G3BP1 condensation in an RNA-containing system and suppresses stress granule accumulation under different stress conditions. (A) Representative fluorescence images of purified GST or GST-G3BP1 (20 μM) incubated with total RNA (100 ng/μL). Proteins were labeled with Alexa Fluor 405, and RNA was stained with PI. Scale bars, 50 μm. (B) Representative fluorescence images of GST-G3BP1 condensates formed in the presence of total RNA and the indicated concentrations of MB (10, 20, or 50 μM). Scale bars, 50 μm. (C) Quantification of condensate number, mean area, and mean fluorescence intensity in (B), normalized to the DMSO control (100%). Data are presented as mean ± SD from three independent LLPS reactions (n = 3). Statistical significance was determined using ordinary one-way ANOVA followed by Dunnett’s multiple-comparisons test against the DMSO control. (D) Representative IF images of G3BP1-positive SGs in CFPAC-1 cells subjected to nutrient deprivation or heat shock with or without MB (10 μM). G3BP1 is shown in red and DAPI-stained nuclei in blue. Scale bar, 10 μm. (E) Quantification of SG accumulation in (D), expressed as the percentage of cellular area occupied by G3BP1-positive puncta. (F) Representative IF images of G3BP1-positive SGs in MIA PaCa-2 cells subjected to nutrient deprivation or heat shock with or without MB (10 μM). Scale bar, 10 μm. (G) Quantification of SG accumulation in (F), using the SG index described in (E). For (E) and (G), data are presented as mean ± SD from three independent biological experiments (n = 3). Statistical significance was determined using ordinary one-way ANOVA followed by Tukey’s multiple-comparisons test. ns, not significant.

**Fig. S4.**
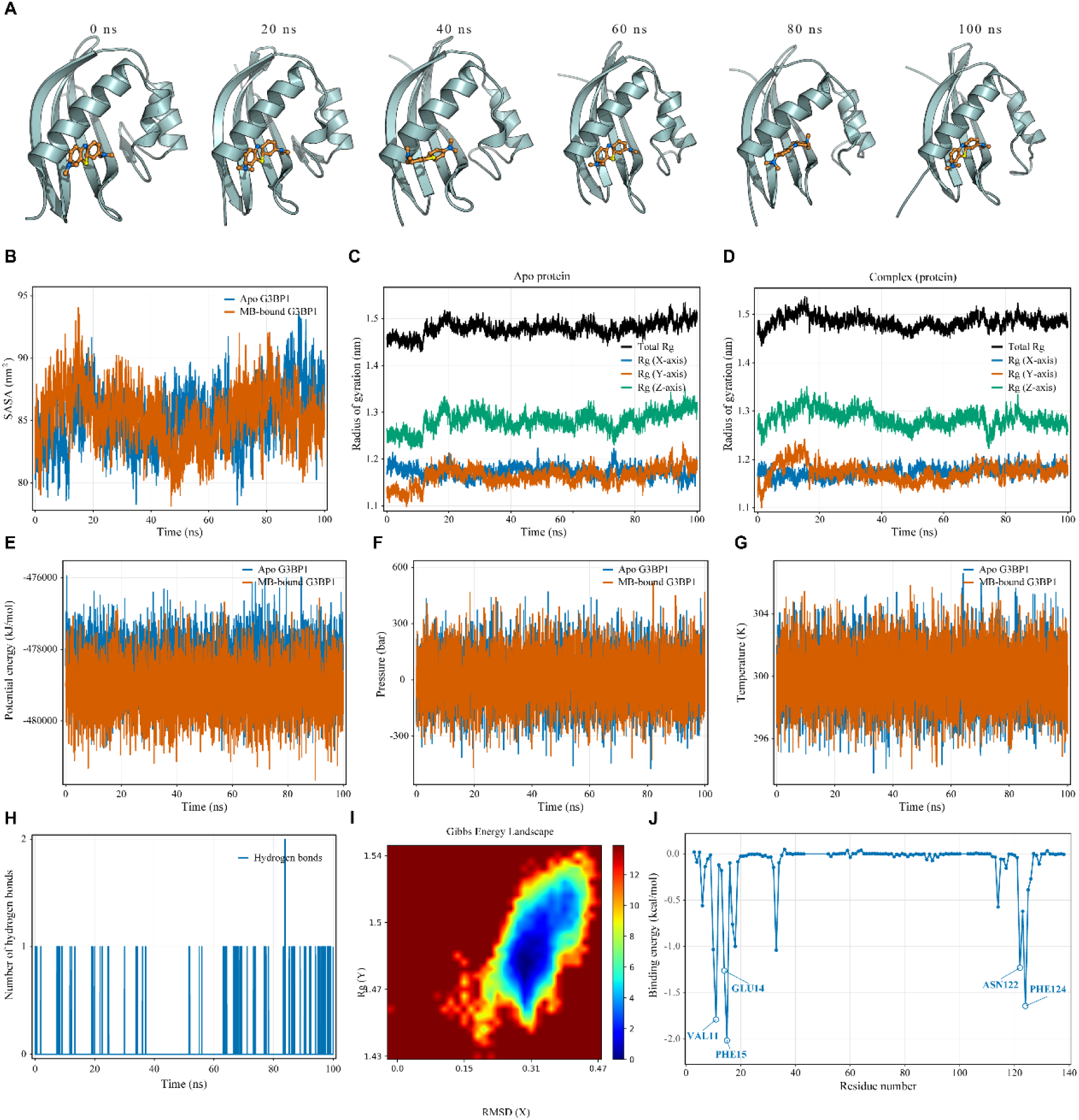
Additional molecular dynamics analyses of the modeled G3BP1– methylene blue complex. (A) Representative conformations of the modeled G3BP1 NTF2L domain–MB complex extracted from the 100-ns MD trajectory at 0, 20, 40, 60, 80, and 100 ns. (B) SASA profiles of apo G3BP1 and MB-bound G3BP1 during the 100-ns MD simulations. (C and D) Total Rg and its X-, Y-, and Z-axis components for apo G3BP1 (C) and MB-bound G3BP1 (D). (E–G) Time-dependent profiles of potential energy (E), pressure (F), and temperature (G) for the apo G3BP1 and G3BP1–MB simulation systems. (H) Number of intermolecular hydrogen bonds between G3BP1 and MB throughout the 100-ns MD simulation. (I) Two-dimensional Gibbs free-energy landscape of the modeled G3BP1–MB complex projected onto RMSD and Rg. (J) MM/PBSA per-residue energy decomposition of the modeled G3BP1–MB complex, showing the calculated contributions of individual G3BP1 residues to the interaction energy. Selected residues with favorable energy contributions are highlighted. The analysis was performed using trajectory frames from the final 10 ns (90–100 ns) of the simulation.

**Fig. S5.**
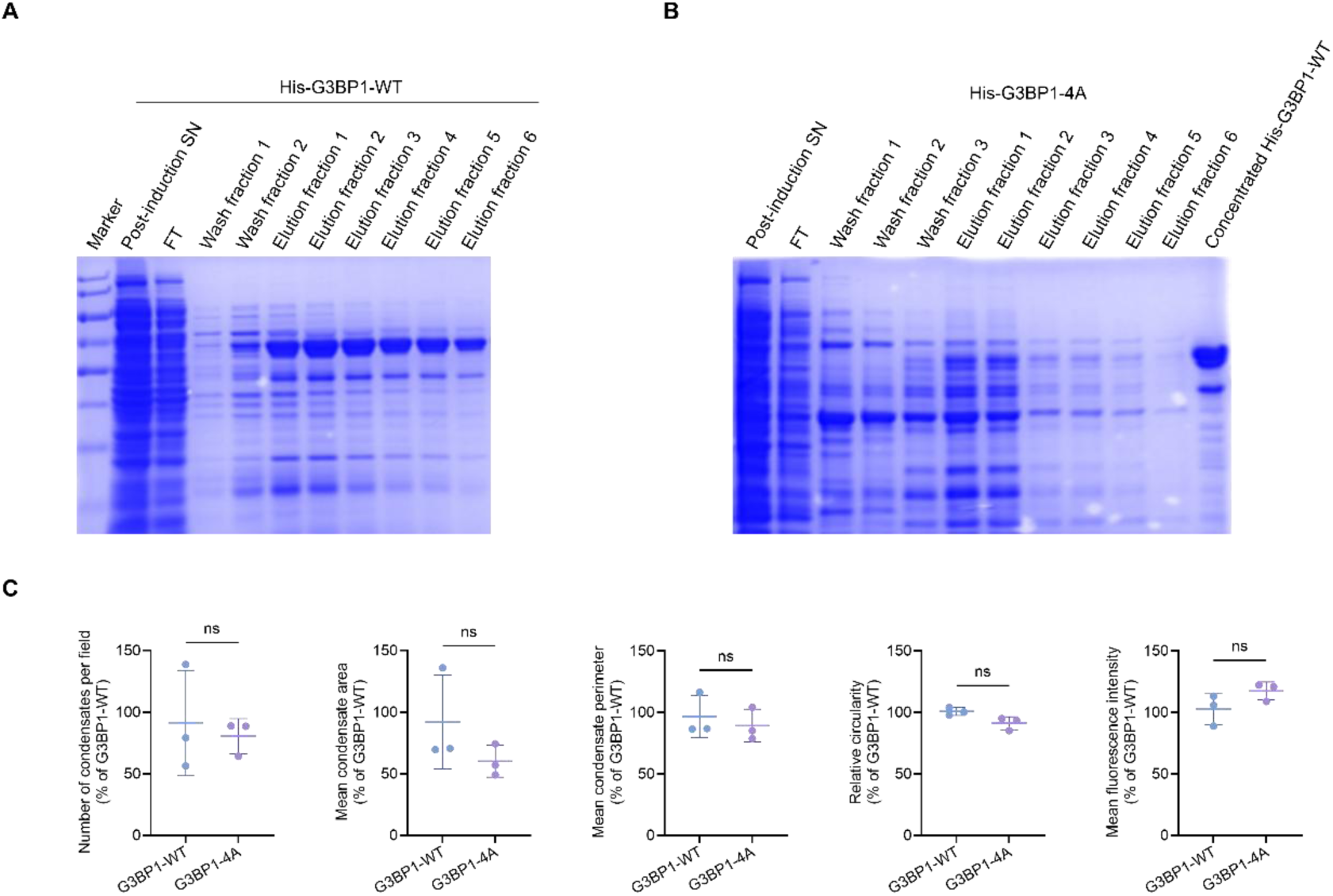
Purification of recombinant G3BP1 proteins and comparison of the basal condensation properties of G3BP1-WT and G3BP1-4A. (A) SDS-PAGE analysis of His-G3BP1-WT during affinity purification, showing the post-induction bacterial supernatant, flow-through (FT), wash fractions, and sequential elution fractions. (B) SDS-PAGE analysis of His-tagged G3BP1-4A during affinity purification, showing the post-induction bacterial supernatant, FT, wash fractions, and sequential elution fractions. Concentrated His-G3BP1-WT was included in the final lane as a reference. (C) Comparison of the basal condensation properties of purified His-G3BP1-WT and His-G3BP1-4A in a reconstituted system without added RNA containing 50 μM protein and 20% (w/v) Ficoll 400 in the absence of MB. Condensate number, mean area, mean perimeter, circularity, and mean fluorescence intensity were quantified using Fiji/ImageJ. Values were normalized to G3BP1-WT (100%). Data are presented as mean ± SD from three independent LLPS reactions (n = 3). Statistical significance was determined using an unpaired two-tailed Welch’s t-test.

**Fig. S6.**
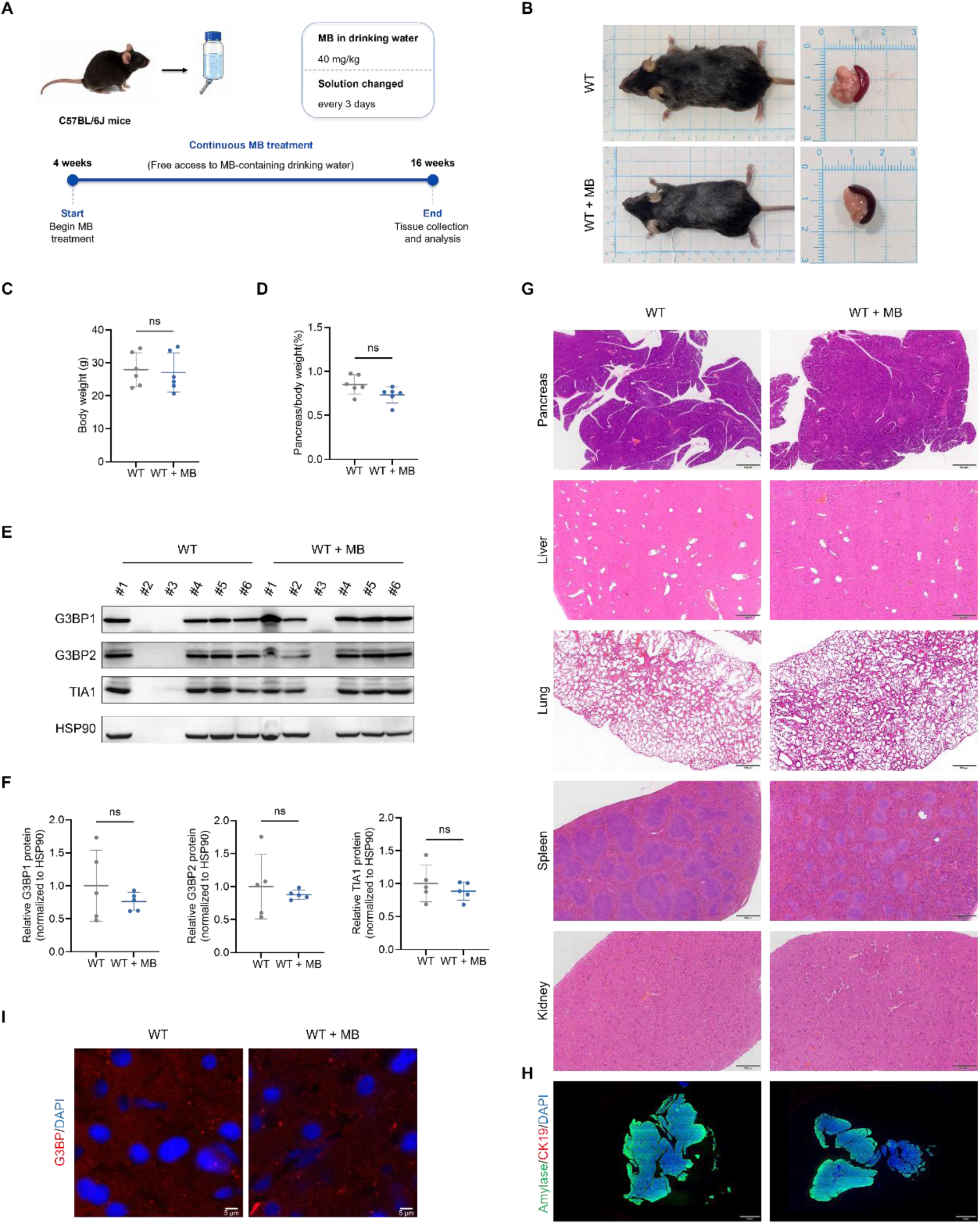
Higher-dose methylene blue administration is well tolerated in wild-type mice under the tested regimen. (A) Schematic overview of the MB administration regimen. Four-week-old WT C57BL/6J mice received MB in drinking water at a nominal dose of 40 mg/kg for 3 months. MB-containing drinking water was replaced every 3 days. (B) Representative photographs of WT mice and excised pancreata from vehicle-and MB-treated groups. (C) Body weights of WT mice at the experimental endpoint. (D) Pancreas-to-body weight ratios of WT mice at the experimental endpoint. (E) Representative immunoblots showing G3BP1, G3BP2, and TIA1 protein abundance in pancreatic tissues from vehicle- and MB-treated WT mice. HSP90 served as the loading control. (F) Densitometric quantification of G3BP1, G3BP2, and TIA1 protein abundance in (E), normalized to HSP90. One sample per group was excluded from densitometric analysis because of protein degradation. (G) Representative H&E-stained sections of the pancreas, liver, lung, spleen, and kidney from vehicle- and MB-treated WT mice. Scale bars, 500 μm. (H) Representative pancreatic tissue IF images showing amylase (green), CK19 (red), and DAPI (blue). Scale bars, 1 mm. (I) Representative pancreatic tissue IF images showing G3BP1 (red) and DAPI-stained nuclei (blue). Scale bars, 5 μm. Data are presented as mean ± SD. For (C) and (D), n = 6 mice per group; for (F), n = 5 mice per group. Statistical significance was determined using an unpaired two-tailed Welch’s t-test.

**Fig. S7.**
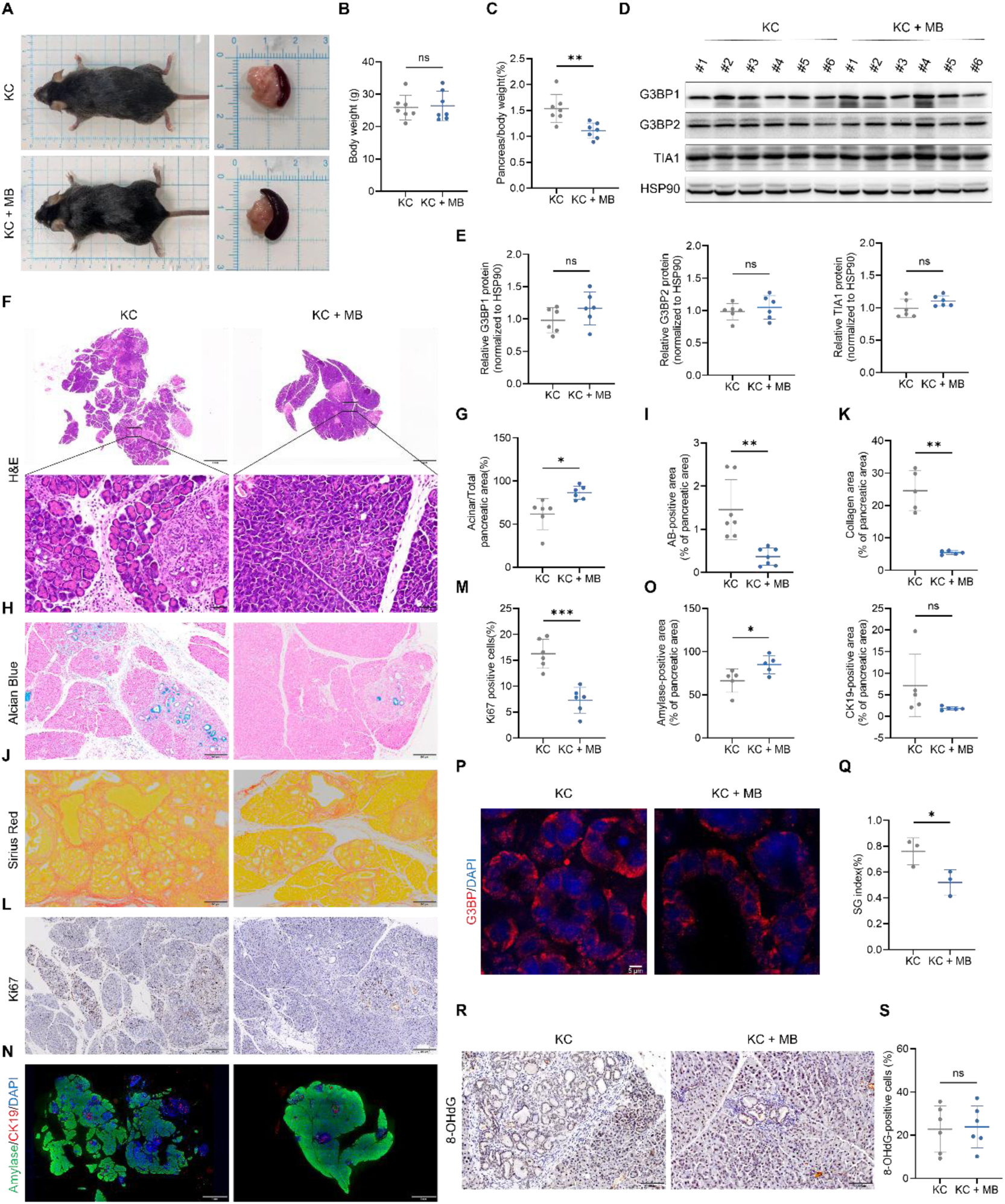
Chronic high-dose methylene blue administration attenuates pancreatic preneoplastic lesions in *Kras*-driven KC mice. (A) Representative photographs of KC mice and excised pancreata following administration of vehicle or MB in drinking water at a nominal dose of 40 mg/kg for 3 months. (B) Body weights of vehicle- and MB-treated KC mice at the experimental endpoint. (C) Pancreas-to-body weight ratios of vehicle- and MB-treated KC mice. (D) Representative immunoblots showing G3BP1, G3BP2, and TIA1 protein abundance in pancreatic tissues. HSP90 served as the loading control. (E) Densitometric quantification of G3BP1, G3BP2, and TIA1 protein abundance in (D), normalized to HSP90. (F) Representative H&E-stained pancreatic sections and enlarged views of the indicated regions. Scale bars, 1 mm (overview) and 50 μm (enlarged views). (G) Quantification of preserved acinar tissue area as a percentage of the total analyzed pancreatic tissue area. (H) Representative Alcian Blue-stained pancreatic sections showing mucin-positive ductal lesions. Scale bars, 200 μm. (I) Quantification of Alcian Blue-positive lesion area as a percentage of the total analyzed pancreatic tissue area. (J) Representative Sirius Red-stained pancreatic sections showing collagen deposition.Scale bars, 200 μm. (K) Quantification of the Sirius Red-stained area fraction, expressed as the percentage of red-stained area relative to the combined red- and yellow-stained areas. (L) Representative Ki-67 immunohistochemical staining of pancreatic tissues. Scale bars, 200 μm. (M) Quantification of Ki-67-positive cells as a percentage of the total evaluated cells. (N) Representative pancreatic tissue IF images showing amylase (green), CK19 (red), and DAPI (blue). Scale bars, 1 mm. (O) Quantification of amylase-positive acinar area and CK19-positive ductal area, each expressed as a percentage of the analyzed pancreatic tissue area. (P) Representative confocal IF images showing G3BP1-positive puncta (red) and DAPI-stained nuclei (blue) in pancreatic tissues. Scale bars, 5 μm. (Q) Quantification of the pancreatic SG index, defined as the percentage of analyzed pancreatic tissue area occupied by G3BP1-positive puncta. (R) Representative immunohistochemical staining of 8-OHdG in pancreatic tissues. Scale bars, 100 μm. (S) Quantification of 8-OHdG-positive cells as a percentage of the total evaluated cells. Quantitative data are presented as mean ± SD. The numbers of mice analyzed per group were as follows: n = 7 for (B), (C), and (I); n = 6 for (E), (G), (M), and (S); n = 5 for (K) and (O); and n = 3 for (Q). Histological area measurements were obtained from whole-section images, whereas three to five confocal fields were analyzed per mouse for tissue IF. Each mouse was considered an independent biological replicate. Statistical significance was determined using an unpaired two-tailed Welch’s t-test.

